# Structural and Mechanistic Basis of FapB Accessory Functional Amyloid Assembly in *Pseudomonas*

**DOI:** 10.64898/2026.10.01.755530

**Authors:** Abdulkadir Tunc, Chang-Hyeock Byeon, Jan-Hannes Schäfer, Kasper Holst Hansen, Danielle A. Grotjahn, Maria Andreasen, Md Rejaul Hoq, James F. Conway, Ümit Akbey

## Abstract

Functional amyloid formation in *Pseudomonas* biofilms is coordinated by the six-gene Fap operon (*fapA-fapF*), which governs the production, secretion, and assembly of extracellular fibrils that reinforce the biofilm matrix. FapC, FapB, and FapE constitute the extracellular amyloid. FapC is the major fibrillar component, whereas FapB and FapE occur at lower abundance and other accessory Fap proteins regulate fibril composition, processing, and secretion. We recently determined the structure of FapC, a cross-β-solenoid major *Pseudomonas* extracellular fibril. The assembly mechanism and architecture of FapB, however, have remained undefined. Here, we elucidate the structural and mechanistic basis of FapB accessory amyloid, from an intrinsically disordered monomer to a cross-β-solenoid fibril. Using well-defined monomeric preparations, controlled fibrillization assays, NMR spectroscopy, cryo-EM, and structure prediction, we show that FapB efficiently self-assembles into structurally homogeneous helical fibrils. FapB fibrillization is highly sensitive to its initial oligomeric state and purity, as well as to environmental factors such as pH. Monomeric FapC does not measurably perturb the disordered conformational ensemble of FapB, whereas preformed FapB and FapC fibrils reciprocally cross-seed each other, revealing direct kinetic coupling between the two amyloid components. Finally, we present a 3.4 Å resolution cryo-EM structure of the FapB accessory fibril, revealing a compact cross-β-solenoid architecture closely related to that of FapC. Together, these findings establish FapB as an independently amyloidogenic, structurally defined component of the Fap system, provide structural basis for the first of accessory cross-β-solenoid fibril, and reveal a mechanistic framework for the post-secretion assembly of multicomponent extracellular functional amyloids in *Pseudomonas* biofilms.

## INTRODUCTION

Protein aggregation and amyloid formation have historically been associated predominantly with neurodegenerative disorders. (*1, 2*) Over the past two decades, this perspective has shifted with the recognition that amyloid structures can serve beneficial and tightly regulated biological functions.(*3, 4*) These so-called functional amyloids are now known to occur across all domains of life, supporting a wide spectrum of physiological processes, from virulence and biofilm formation in bacteria to peptide hormone storage, pigmentation, memory formation, and programmed cell death.(*5–8*) The rapidly expanding study of functional amyloids has established amyloid formation as a broadly employed biological strategy rather than exclusively a pathological process.(*4, 9, 10*)

A defining feature that distinguishes functional amyloids from pathological aggregates is the degree of regulatory control, while pathologic amyloids are uncontrolled anomalies. Functional amyloids are produced through genetically encoded pathways that coordinate expression, secretion, and assembly, ensuring that fibril formation occurs at the appropriate location, time, and rate. Such regulation is essential because amyloid fibrils are among the most stable protein assemblies known, and their uncontrolled formation can compromise cellular integrity and function. While high-resolution structures of mature functional amyloid fibrils involved in bacterial biofilm extracellular matrix (ECM) are increasingly available,(*4, 11–17*) the molecular principles underlying their tightly regulated assembly, particularly during the early post-secretion stages, remain poorly defined.

In *Pseudomonas* species, extracellular functional amyloid formation in biofilm is mediated by the Fap (functional amyloid in Pseudomonas) operon, which encodes six proteins (FapA-FapF) that coordinate secretion, assembly, and fibril maturation (**Fig. 1A**).(*18, 19*) The resulting fibrils reinforce the biofilm ECM, providing protease resistance, mechanical strength, and hydration retention. These properties help biofilms withstand mechanical stress, desiccation, and mediate antimicrobial resistance (AMR). Among the Fap proteins, FapB, FapC, and FapE contribute directly to ECM formation as secreted extracellular amyloids.(*19*) FapA, FapD, and FapF regulate distinct stages of the pathway: FapA modulates FapC fibrillation and fibril composition in the periplasm, FapD is a C39-family protease involved in precursor processing in the periplasm, and FapF forms a β-barrel outer-membrane channel that exports amyloidogenic subunits as intrinsically disordered protein (IDP) monomers into the extracellular environment.(*18–22*) Once secreted, the amyloidogenic components undergo coordinated conformational transitions from IDP monomeric precursor states into their mature extracellular amyloid folds.

**Fig. 1:**
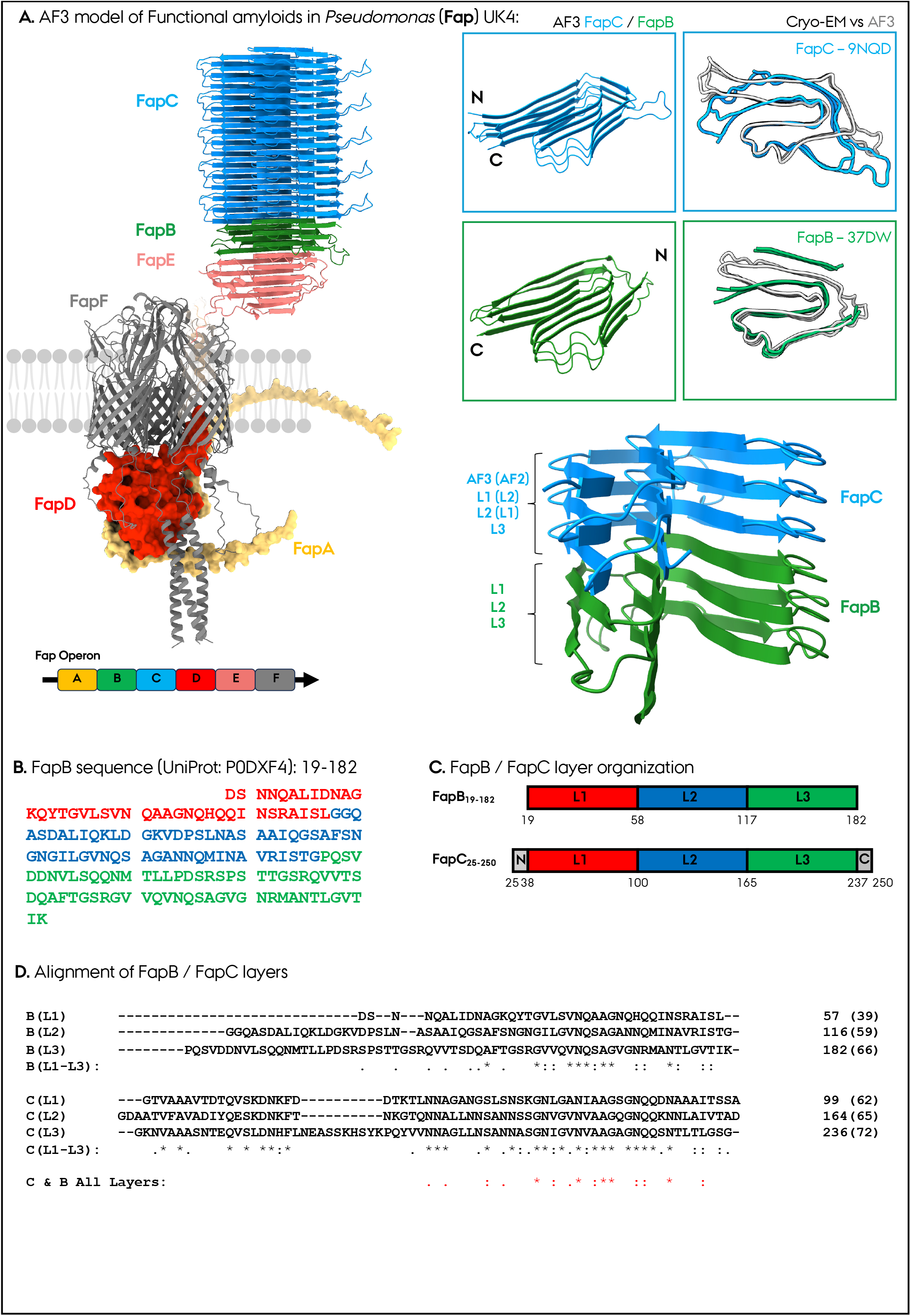
Structural and modular organization properties of functional amyloids in Pseudomonas (Fap). **A.** AF3 modeling of FapA-F proteins and operon structure. The insert shows a close-up view of FapB (green) and FapC (blue), and the fold similarities/differences of the AF3 models (grey) with the Cryo-EM structures (green and blue) determined by us in this and previous work. The side view of FapB and FapC shows the cross-β-solenoid folds, each composed of three independent layers. These layers organize differently in AF2/AF3 models for FapC and are depicted. **B.** The sequence of the FapB functional amyloid protein, color-coded following the domain organization shown in **C.** for FapB and FapC. **D.** Multiple sequence alignment (MSA) of six FapB and FapC layers (three from each) treated as individual proteins. The MSA conservation is shown for FapB and FapC separately below their sequences, given along with the total conservation of all six layers at the bottom.

Like the CsgA/CsgB system of *Escherichia coli* and TasA/TapA from *B. subtilis*,(*23–28*) the FapC/FapB system provides a powerful model for investigating how bacteria control extracellular amyloid formation while avoiding premature intracellular aggregation. Among these, TasA/TapA are not cross-β-solenoid protein fibrils and have globular β-sheet rich protein folds,(*12, 29*) while both the CsgA/CsgB and the FapC/FapB are predicted/shown to have or shown to have a cross-β-solenoid structure.(*30*) Encoded by a single operon, the Fap machinery is broadly conserved across *Pseudomonas* species.(*19*) In the clinically important pathogen *P. aeruginosa*,(*31, 32*) Fap amyloids contribute to persistent biofilms associated with chronic infections, including those in airways of cystic fibrosis, where ECM stability promotes disease persistence and antibiotic tolerance.(*31, 32*)

Assembly within the Fap system involves distinct but interconnected roles for its extracellular amyloid-forming components. FapB, FapC, and FapE occur at different stoichiometries within mature ECM fibrils.(*19*) FapC forms the dominant fibrillar scaffold, whereas FapE is consistently detected in mature fibrils and may contribute to fibril anchoring or assembly control. FapB is generally regarded as an accessory component and has been proposed to nucleate FapC fibrillation, by analogy to the CsgA/CsgB or TasA/TapA.(*23–28*) However, this proposed nucleator function has largely been inferred rather than directly demonstrated. Despite intensive previous work on FapC and FapA biophysics,(*4, 18, 19, 21, 33–41*) mechanisms and structures, investigations on accessory amyloid FapB are limited.(*42*) Moreover, no accessory cross-β amyloid structure is available yet.

Several observations suggest that the role of FapB is more complex than that of a minor structural component. FapB displays sequence features, strain-dependent variation, and aggregation behavior distinct from FapC. (*18–20, 43, 44*) Although extracellular Fap amyloid fibrils from *P. aeruginosa* and *P. fluorescens* are composed predominantly of FapC, smaller amounts of FapB and FapE are incorporated into the mature fibrils. Disruption of FapA markedly alters this composition without preventing amyloid formation, producing fibrils composed predominantly of FapB with only a minor FapC fraction.(*19*) This indicates that FapA regulates or facilitates the relative incorporation of FapB and FapC into the fibril. In contrast, FapE is a low-abundance fibril-associated component whose deletion largely abolishes extracellular amyloid production, supporting an essential role in Fap biogenesis or secretion rather than as a major structural subunit.(*19*) Together, these findings suggest that FapB is an accessory fibril component with a tunable, condition-dependent role in Fap assembly.

A major limitation in prior work is the lack of mechanistic studies using well-defined, monomeric FapB. Fibrillation kinetics are highly sensitive to the initial oligomeric state, trace quantities of pre-existing aggregates, and sample purity, yet most existing studies have relied on affinity-purified preparations in which these variables are difficult to isolate and control.(*40*) This raises questions about whether previously observed fibrillation behavior reflects the intrinsic biochemical properties of FapB or preparation-dependent effects. Moreover, the structural and mechanistic interactions of FapB with FapC and FapE during the early stages of extracellular assembly remain poorly understood.

To address these questions, we established a mechanistic framework for studying the extracellular functional amyloid FapB from *Pseudomonas fluorescens* UK4 using a well-defined, highly pure monomeric starting state. Using optimized size-exclusion chromatography (SEC), we isolate strictly monomeric FapB and demonstrate that its fibrillation behavior differs markedly from that of immobilized metal ion affinity chromatography (Ni-NTA) purified preparations of His-tagged proteins despite their comparable nominal purity. We integrate controlled fibrillation and seeding assays, negative-stain and cryo-electron microscopy (Cryo-EM), solution NMR spectroscopy, AlphaFold (AF)-based modeling, and molecular dynamics simulations to determine how FapB assembles independently and cooperatively with FapC. Our results establish FapB as a regulated accessory amyloid whose assembly depends on its oligomeric state, sample purity, the surrounding ionic environment, and the availability of interacting amyloid partners. Structural modeling and kinetic analyses support a model where FapB and FapC are secreted as unfolded (IDP) monomers that are weakly interacting, which subsequently assemble through cooperative extracellular interactions. This study establishes the role of FapB as an effective accessory nucleator fibril and reveals broader principles through which bacteria construct robust, yet adaptable multicomponent amyloid matrices. These mechanistic insights may ultimately reveal new opportunities for disrupting amyloid-supported biofilms.

## RESULTS

### Predicted organization of extracellular Fap functional amyloids in the *Pseudomonas* biofilm ECM

Extracellular functional amyloid formation in *Pseudomonas* biofilms is mediated primarily by FapB, FapC, and FapE (**Fig. 1A**). These proteins are exported through the FapF outer-membrane pore as unfolded monomers, undergoing a controlled transition into their mature amyloid folds.(*21, 22, 41*) We recently determined FapC’s conformational transition from an unfolded monomer into a folded AF2-like monomer, which then reorganizes into a cross-β-solenoid L2-L1-L3 amyloid fibril architecture via NMR, cryo-EM and MD simulations.(*17*) To date, there are no high-resolution structures for FapB and FapE.

To explore the potential structures and interactions of Fap members, we used AF to model FapB individually and together with FapC and FapE across stoichiometries (**Fig. SI1**). Both FapB and FapC were predicted to adopt β-sheet-rich, multilayer cross-β-solenoid folds with repeating strand-loop motifs: FapB in L1-L2-L3 and FapC in either L1-L2-L3 (AF3) or L2-L1-L3 (AF2) (**Fig. SI2**). FapE’s AF3 model is β-sheet-rich but does not form a complete β-solenoid (**Fig. 1A**). AF3 models consistently predicted a conserved FapE → FapB → FapC spatial complex organization, where FapE’s spatial terminal region consistently interfaces with bottom L3 layer of both FapB and FapC, and the top layer of FapE (residues 102–138) align with L3 of both FapB and FapC (**Fig. SI1**). These support a higher-order Fap complex in which membrane-anchored FapE recruits FapB, FapB bridges FapE and FapC and templates FapC fibrillation. While these models don’t establish kinetics or directionality, they provide a structural hypothesis we test in later sections by kinetic assays on FapB and FapC.

We next analyzed FapB and FapC’s sequence and structural predictions. The AF3 model of FapB (UniProt ID: P0DXF4 (previously C4IN69); AF-P0DXF4) adopts an ordered three-layer L1–L2–L3 β-solenoid fold resembling FapC (UniProt ID: P0DXF5 (previously C4IN70); PDB: 9NQD, 9U4U), in particular at the S-shaped core regions (**Fig. 1A**).(*17*) However, the confidence scores of FapB structure predictions were lower in the N-/C-terminal regions and in residues 58–77 and 116–144 constituting the irregular/asymmetric β-sheet region, while the S-shaped core region (22–57, 78–115, 145–181) has high confidence (**Fig. SI2**). Similarly, the FapC AF3 model has low confidence at the flexible loop regions and cross-over areas between layers. FapB exhibiting lower confidence and asymmetric β-sheet organization may indicate higher conformational plasticity, potentially relevant for inter-subunit interactions with itself or FapC. Despite only ∼30% sequence identity (**Fig. SI3**), AF3 predictions indicate that the FapB and FapC share very similar modular β-solenoid architectures (**Fig. 1A-C**). FapC’s three layers each contain five β-strands, whereas FapB’s L2 and L3 have five β-strands and L1 has only four. This results in four β-sheets containing three strands and a fifth β-sheet containing only two, producing an asymmetric solenoid fold with a structural gap one side of the β-solenoid (**Fig. 1A**). Sequence alignment of FapB and FapC reveals that FapC is longer at the L1 and termini and FapB is missing a β-strand at L1, which is the reason for the irregular/asymmetric β-sheet region and shorter repeats of FapB compared to FapC (**Fig. 1D**, **SI3**). Both FapB and FapC contain three imperfect repeats predicted by RADAR, (**Fig. SI3**). FapC’s repeats (∼60, 63, 67 residues for L1-L3, respectively) are longer and separated by flexible turns, while FapB has shorter repeats (35, 39, 41 residues for L1-L3, respectively) again due to the shorter L1 section (**Fig. SI3, SI4**). FapB’s shorter, more asymmetric L1 displays greater irregularity and may reduce its self-nucleation potential and subsequent growth into a mature fibril *in vivo*. Since FapB and FapC solenoids both have a spatial third-dimension component to their structures due to the β-solenoid fold, we performed a modified sequence alignment protocol (**Fig. 1D**). We separated the individual six layers of both FapB and FapC and performed sequence alignment of the resulting peptide sections as if they were separate protein sequences. As a result, we obtained a less spatially constrained fold alignment and performed an analysis similar to the original repeat prediction by RADAR but not constrained to one protein sequence and comprising all six layers. This method revealed the sequence alignment of FapB and FapC layers and indicated the regions/residues of importance for β-solenoid structural matching of FapB with FapC.

Finally, sequence comparisons among different *Pseudomonas* strains highlight strain-specific variation in FapB (**Fig. SI5**). For instance, *P. fluorescens* UK4 FapB (182 aa, UniProt ID: P0DXF4) slightly differs in length and amino acid composition from *P. aeruginosa* PAO1 (189 aa, UniProt ID: Q9I2E9) and *P. fluorescens* Pf0-1 (189 aa, Pseudomonas Genome Database: Pfl01_2597), especially in loop and linker regions mapping to the L3 layer (**Fig. SI5**). Overall, the structural fold is very similar among these, except for small changes in the hydrophobicity and electrostatic surface when modelled in complex with FapC (modelled in a complex with 1x FapB + 5x FapC, with the corresponding FapC from the same strain) (**Fig. SI5**). We selected the mature form of secreted FapB from the UK4 strain (residues 19-182, excluding the signal peptide) as the representative construct for all downstream experiments, as a follow-up to our previous FapC cryo-EM structure from the UK4 strain.

### Monomer, oligomeric state and purity influence FapB fibrillation

The position of the His-tag (N-vs. C-terminal) significantly influenced *P. fluorescens* UK4 FapB (UniProt ID: P0DXF4) fibrillation. This was similarly observed for prion proteins.(*45*) To evaluate this effect, we generated two FapB constructs with either an N- or C-terminal His-tag (**Fig. SI3**). 40 μM of both variants formed fibrils under identical conditions but exhibited markedly different kinetics (T_lag_ & T_50_) and maximum ThioflavinT (ThT) fluorescence intensities (ThT_max_) (**Fig. SI6**). The N-terminal construct showed a prolonged lag phase (T_lag_ > 10 h) and lower ThT_max_, while the C-terminal variant fibrillated more rapidly (T_lag_ ∼1 h) with higher ThT_max_. These trends remained consistent across tested concentrations. Based on these results, we selected the C-terminally tagged FapB for all subsequent experiments.

FapB fibrillation was strongly affected by the initial oligomeric state, similar to our previous results for FapC.(*40*) To evaluate its intrinsic assembly properties, we compared the fibrillation kinetics of FapB purified by Ni-NTA affinity chromatography with those of FapB subjected to additional SEC. Although Ni-NTA purification yielded high purity (>95%) by SDS-PAGE, concentrated samples contained high-molecular-weight (HMW) oligomers and minor low-molecular-weight (LMW) species (**Fig. 2A**, **SI6**). SEC purification enabled the isolation of a well-defined monomeric FapB fraction (**Fig. 2A**, **SI6**), as supported by its SEC elution profile, SDS-PAGE analysis and mass spectrometry, which confirmed the identity and high purity of the sample.

**Fig. 2:**
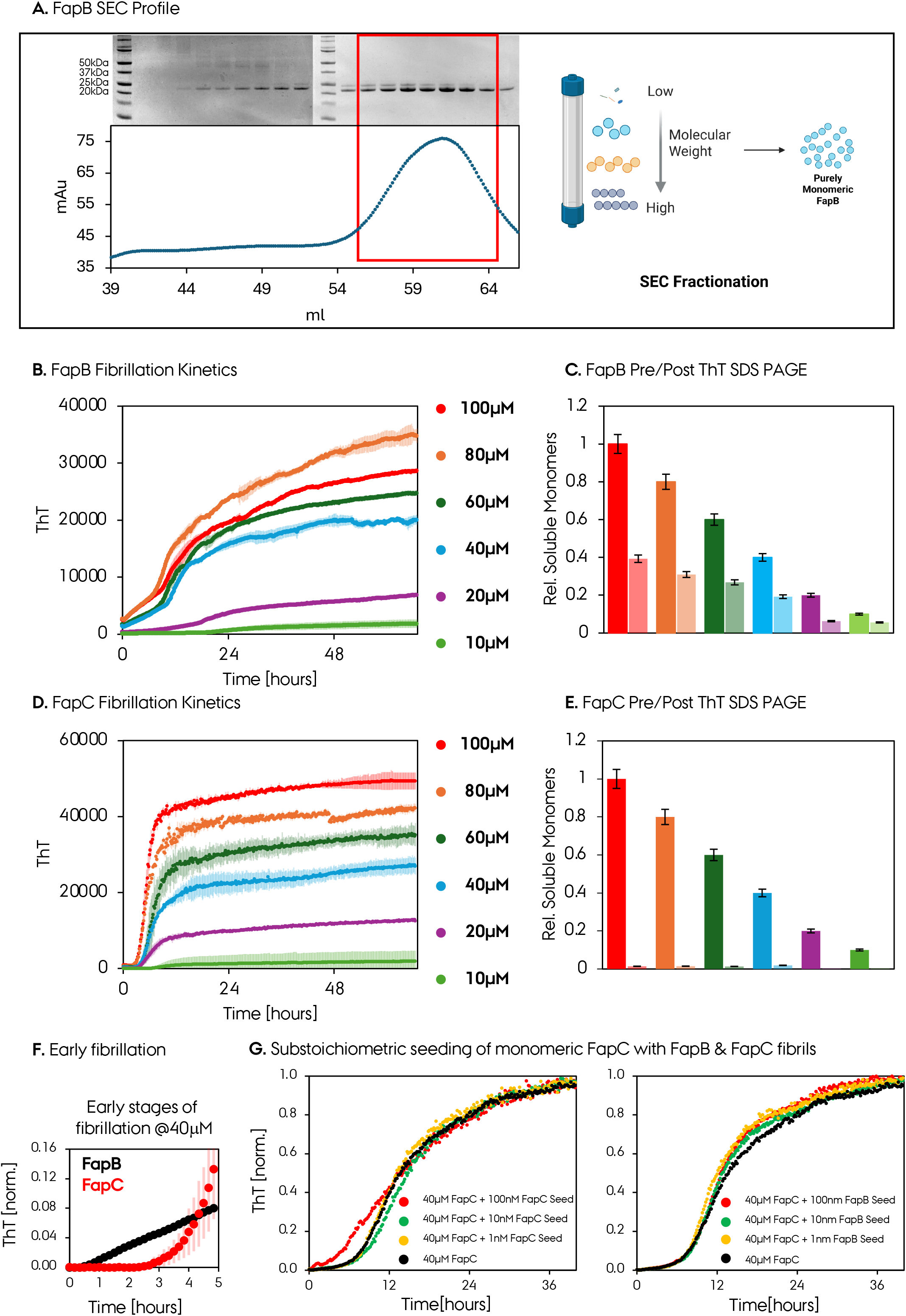
High-purity SEC-purified FapB and FapC and their fibril formation properties. **A.** SEC separates impurities and oligomeric states of FapB into defined impure and pure monomeric and non-monomeric pools, enabling controlled assessment of their impact on fibrillation behavior. SEC elution profiles, the collected fractions (in red), and their SDS-PAGE profiles are shown. **B,D.** The fibrillation kinetics and extent were quantified for freshly desalted pure monomeric FapB and FapC samples at different concentrations at pH 7.8. **C,E.** The extent of fibrillation is quantified by SDS page pre/post ThT assay for each concentration. **F.** The comparison of the early fibrillation steps of FapB and FapC. FapB has a much smaller Tlag and starts fibrillation much faster compared to FapC. The effect of substoichiometric FapB and FapC seeds on pure monomeric FapC. 40 μM FapC was incubated with/out 1, 10, and 100 nM seeds, respectively. The substoichiometric seeding did not change the overall ThT curve, indicating a lack of secondary nucleation.

Specifically, SEC-purified FapB produced higher ThT_max_ values and faster fibrillation than Ni-NTA purified samples (**Fig. SI6**). For C-terminally His-tagged (C-His) FapB, SEC-purified samples exhibited a T_lag_: 1 h, a T_50_: 19 h, and a ThT_max_: ∼15,000, whereas the Ni-NTA-purified C-His FapB sample showed minimal ThT signal after 48 h. The N-terminal His-tagged (N-His) samples showed similar trends, and the SEC-purified FapB displayed improved kinetics (T_lag_: 12 h, T_50_: 32.5 h, ThT_max_: ∼5,500), whereas the Ni-NTA-purified form fibrillated more slowly and weakly (T_lag_: 16 h, T_50_: 36 h, ThT_max_: ∼1,300).

This behavior is consistent with our previous findings with FapC, for which fibrillation was highly sensitive to sample oligomeric heterogeneity.(*40*) In the FapC system, isolated HMW oligomers accelerated fibril formation, while LMW fragments impaired it and monomer-only SEC fractions exhibited longer lag phases.(*40*) Because the Ni-NTA-purified FapB preparations contained both HMW oligomers and LMW species, the present data do not identify which component causes the reduced fibrillation. Instead, they demonstrate that unresolved oligomeric heterogeneity can substantially alter the observed FapB fibrillation kinetics. Together, these results highlight the importance of using strictly monomeric, well-defined protein preparations to study amyloid fibrillation. Trace oligomers or fragments in affinity-purified samples can alter kinetics, obscure mechanistic interpretation, and misrepresent intrinsic behavior despite high apparent purity by SDS-PAGE. FapB, like FapC, is highly sensitive to oligomeric heterogeneity and requires rigorous sample definition to ensure reliable kinetic and structural analysis.

### Monomeric FapB and FapC exhibit distinct fibrillation kinetics

To directly compare their intrinsic fibrillation behavior, SEC-purified monomeric FapB and FapC were analyzed under identical conditions.(*34*) ThT assays showed that C-terminally His-tagged FapB undergoes robust, concentration-dependent fibrillation with sigmoidal kinetics (**Fig. 2B**). The lag phase kinetics are sloped and not flat as expected in a sigmoidal curve, and as a result represent a shorter T_lag_ compared to a near-zero intensity curve observed at the initial time points for FapC. Negative-stain EM and Cryo-EM 2D class averages confirmed the formation of characteristic amyloid fibrils with identifiable cross-beta stacks and a layer line at ∼1/4.8 Å (**Fig. SI7**). FapB amyloids assemble into ∼7.2 nm wide fibrils, approximately 1 nm narrower than FapC fibrils, and seem to be shorter compared to long FapC filaments (**Fig. SI7**). These results demonstrate that monomeric FapB is intrinsically capable of assembling into amyloid fibrils without requiring detectable pre-existing oligomers.

FapB and FapC displayed markedly different kinetic profiles (**Fig. 2B-F**). FapB exhibited a shorter apparent lag phase than FapC (T_lag_ ∼1 h versus ∼4 h), indicating earlier onset of detectable fibrillation. However, FapB reached half-maximal fluorescence later than FapC (T_50_ ∼13 h versus ∼8 h) and displayed a lower final ThT fluorescence (ThT_max_ ∼20,000 versus ∼27,000). Examination of the early time points further emphasizes the shorter lag phase of FapB, due to its non-flat initial ThT curve (**Fig. 2F**). Thus, although FapB initiates detectable assembly earlier, its subsequent fibril growth is slower, whereas FapC undergoes a more rapid transition from nucleation to fibril accumulation.

SDS-PAGE analysis was consistent with these kinetic differences, showing approximately ∼60% depletion of soluble FapB compared with near-complete depletion of FapC (**Fig. 2C,E**). Because ThT fluorescence depends on fibril architecture, dye accessibility and affinity of ThT-binding sites, the lower ThT_max_ of FapB cannot be interpreted directly as a proportional reduction in fibril mass. Nevertheless, the substantial residual soluble FapB indicates less efficient monomer incorporation under these conditions. Together, these results suggest that FapB crosses the initial nucleation barrier more readily, whereas FapC subsequently undergoes faster fibril growth and more efficient conversion of soluble monomer into fibrils.

To further investigate the FapC fibrillation mechanism, we performed low-level seeding experiments using preformed FapC or FapB fibrils. Such substoichiometric seeding can be informative for identifying fibril-surface-catalyzed secondary nucleation, as even very low seed concentrations can markedly accelerate aggregation when secondary nucleation contributes strongly to fibril proliferation.(*34*) FapC monomer (40 μM) was therefore supplemented with 1-100 nM FapC or FapB seeds (**Fig. 2G**). Across this concentration range, neither homologous FapC seeds nor heterologous FapB seeds produced a detectable acceleration of fibrillation or shortening of the lag phase. These results argue against a substantial contribution of seed-dependent secondary nucleation under the conditions tested. Consistent with this interpretation, global kinetic analysis of the concentration-dependent FapB and FapC fibrillation data using AmyloFit (**Fig. SI8**) revealed only a weak dependence of aggregation half-time on monomer concentration. The data were adequately described by a primary nucleation–elongation model, while inclusion of more complex secondary-nucleation pathways did not improve the quality of the fits. Together, the seeding experiments and kinetic modeling support a fibrillation mechanism dominated by primary nucleation followed by elongation for both FapC and FapB under these experimental conditions.(*34*) This behavior contrasts with many pathological amyloid systems in which fibril-dependent secondary nucleation can make a major contribution to aggregate proliferation.

### Monomeric FapB and FapC fibrillation is regulated by stoichiometry and pH

To evaluate how environmental context influences the combined fibrillation of FapB and FapC monomers, we investigated fibrillation kinetics across various FapB stoichiometries (FapB:FapC) to approximate different relative protein abundances that may occur in vivo. We additionally tested two biologically relevant pH conditions, 5.0 and 7.8 (**Fig. 3A,B**). The near-neutral to mildly alkaline pH 7.8 condition approximates selected periplasmic or extracellular environments, whereas pH 5.0 mimics acidic microenvironments associated with biofilm maturation.(*46*) Because FapB is present at much lower abundance *in vivo* (estimated FapB ratio of ∼1:10-1:100),(*18, 19*) we tested a wide stoichiometric range from 20:1 to 1:20 FapB:FapC, thereby including conditions in which either FapB or FapC was limiting. This range accounts for potential variations in relative protein abundance arising from differences in expression, processing, and secretion.

**Fig. 3:**
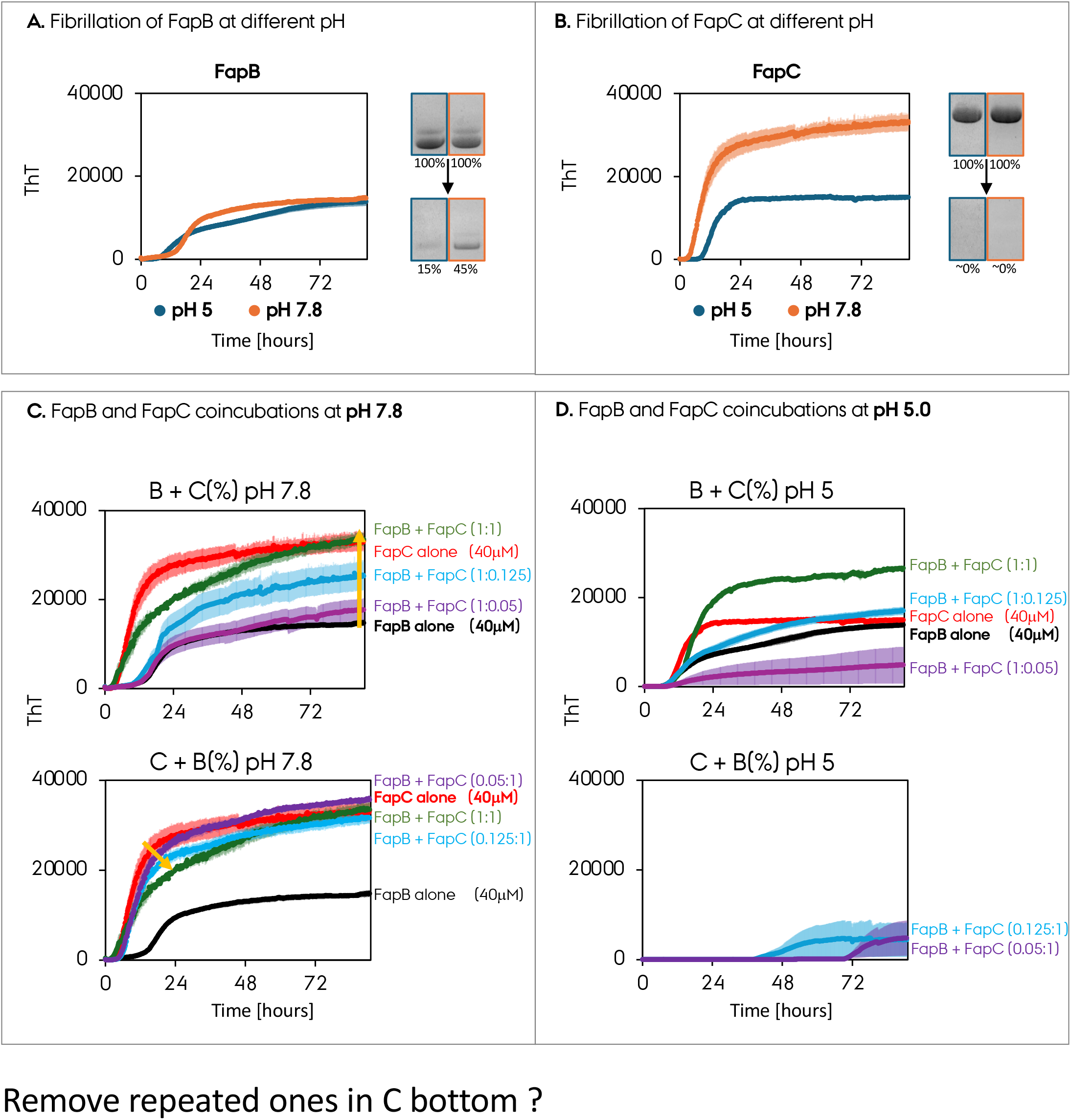
The effect of biologically relevant pH on FapB/FapB fibrillation and co-fibrillation. **A,B.** The ThT fibrillation assay on pure monomeric FapB/FapC at pH 5.0 and 7.8, respectively. The extent of monomer depletion is quantified with SDS-PAGEs. **C,D.** Coincubation of monomeric FapB and FapC at different stoichiometries and pH 5.0 (acidic) or 7.8 (∼neutral). Different stoichiometries are indicated by ratios and color codes.

At pH 7.8, both proteins fibrillated independently but with distinct profiles (**Fig. 3A,B**). FapC formed fibrils with high efficiency (T_lag_: 2.5 h; ThT_max_: ∼30,000) and near-complete monomer depletion by SDS-PAGE. FapB also fibrillated, though less efficiently (T_lag_: 1 h; ThT_max_: ∼15,000), with ∼45% monomer remaining after 90 h. At pH 5.0, FapC still formed fibrils with complete monomer depletion but with slower kinetics (T_lag_: 7 h) and reduced ThT_max_ (∼15,000). In contrast, FapB conversion improved under acidic conditions and although its ThT_max_ remained similar to that at pH 7.8, remaining monomer decreased to ∼15%, and its lag phase increased from approximately 1 h at pH 7.8 to 6 h at pH 5.0. These results show that pH differentially affects FapB and FapC and that ThT intensity does not necessarily scale directly with monomer conversion.

To assess how FapB stoichiometry modulates their combined assembly, we co-incubated FapB and FapC monomers at various ratios (**Fig. 3C,D**). Unlike seeding assays, this setup explores direct monomer-monomer interactions and potential competition or cooperation during nucleation in biofilm ECM. This experiment approximates an extracellular scenario in which FapB and FapC become available for assembly within overlapping time windows. However, ThT kinetics and soluble-protein depletion alone cannot distinguish molecular co-assembly within the same fibrils from the parallel formation of separate FapB and FapC fibrils.

At pH 7.8 (**Fig. 3C**), the combined-assembly landscape was permissive but nonlinear and nonadditive. For instance, the 1:1 FapB mixture yielded a ThT_max_ lower than the sum of the individual components, indicating possible interference or competition at equimolar levels or altered ThT binding in the mixed sample. In FapB-dominant mixtures (top panels), increasing FapC concentration progressively accelerated fibrillation, reducing T_lag_ and increasing ThT_max_ up to approximately the pure FapC level. Notably, when FapB monomer was added at progressively lower ratios relative to FapC monomer (bottom panels), the 1:8 and 1:20 FapB conditions didn’t exhibit a significant change, maintaining signal levels comparable to pure FapC and the 1:1 mixture. This behavior suggests that a threshold abundance of FapC promotes efficient assembly when FapB is present at a low stoichiometric concentration, resembling the proposed relative abundance of the proteins *in vivo*.(*18, 19*) SDS-PAGE confirmed nearly complete monomer depletion of both proteins at these ratios, indicating efficient conversion of both components into insoluble assemblies (**Fig. SI9**).

At pH 5.0, however, this broad compatibility was lost (**Fig. 3D**). Only the 1:1 ratio supported effective fibrillation with an approximately additive ThT_max_. Highly asymmetric conditions (e.g., 20:1 or 1:20) led to substantially prolonged lag phases and diminished ThT signals, with SDS-PAGE showing poor incorporation or depletion of the minority component (**Fig. SI9**). These results indicate that acidic conditions impose stricter constraints on the combined assembly of FapB and FapC. This restriction may result from pH-dependent changes in protonation, intermolecular electrostatics, conformational populations, or fibril-surface compatibility, although the present experiments do not distinguish among these possibilities. In contrast, near-neutral pH facilitates cooperative kinetic behavior across a broader range of FapB ratios.

Altogether, these findings show that FapB-FapC assembly behavior is finely tuned by both stoichiometry and pH. Acidic conditions restrict integration or conversion of the minority species, while near-neutral pH supports broader cooperative behavior, particularly when FapC is abundant. Discrepancies between ThT fluorescence and monomer depletion underscore that pH may affect dye accessibility or fibril morphology. Direct structural characterization will be required to determine whether the mixed preparations produce heteromeric fibrils, separate homomeric fibrils, or a combination of both.

### FapC monomer does not alter the IDP FapB monomer

To determine whether FapC influences the conformational properties of monomeric FapB prior to assembly, we used high-resolution solution NMR to assess potential chemical shift perturbations (CSPs) and signal intensity changes (**Fig. 4**). We acquired 2D ^1^H-^15^N HSQC spectra of uniformly ^15^N-labeled FapB monomer in the presence and absence of unlabeled FapC monomer at a 1:1 molar ratio (**Fig. 4A**). FapB assignments were transferred from previous assignments.(*22*) Both FapB and FapC are intrinsically disordered in their monomeric states, and our previous data show that FapC exhibits local secondary-structure propensities related to elements of its fibrillar fold.

**Fig. 4:**
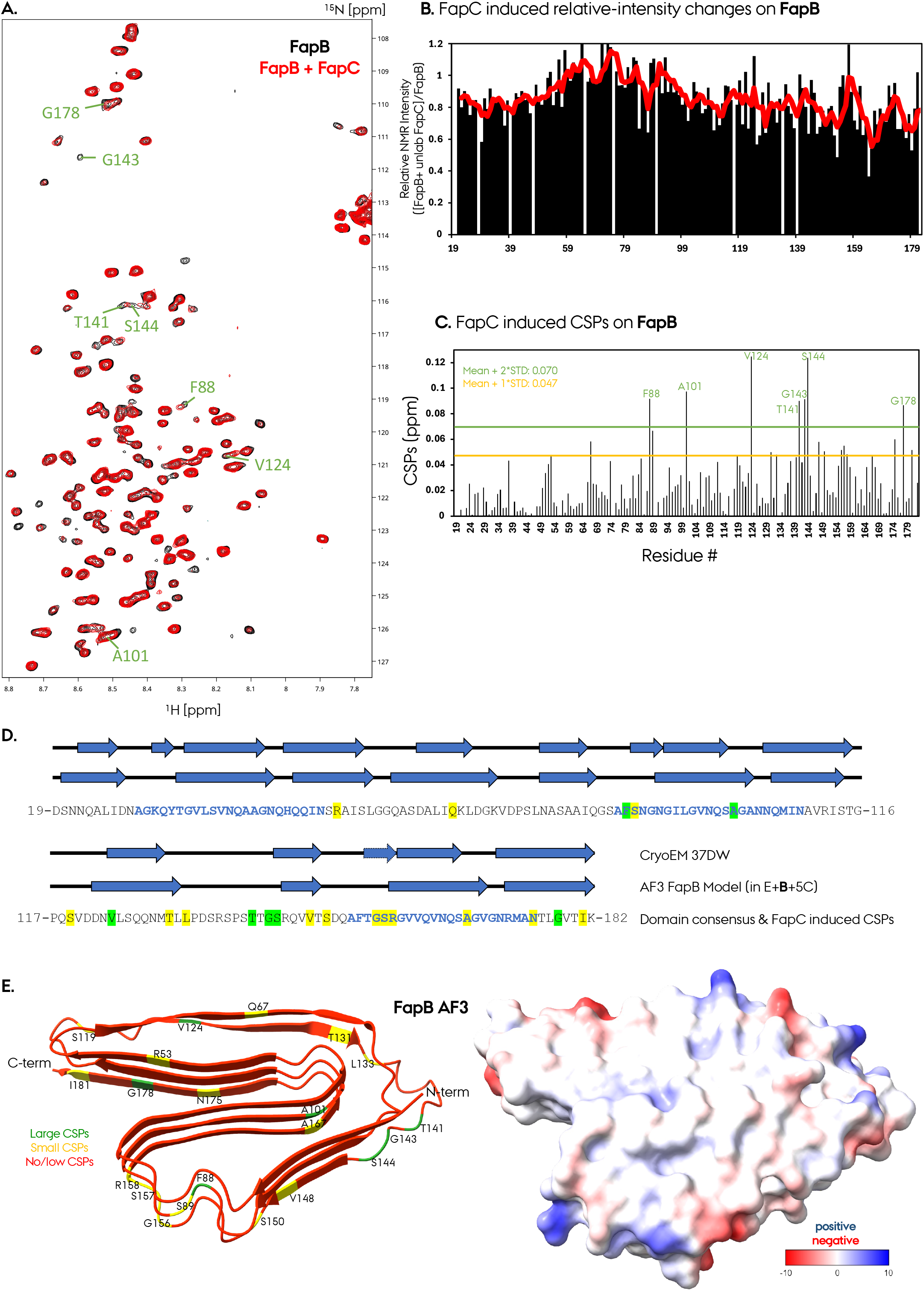
Solution NMR characterization of the effect of monomeric FapC on the monomeric FapB. **A.** 2D ^1^H-^15^N HSQC solution-state NMR spectra of ^15^N isotope-labeled pure FapB (black) and with unlabeled FapC at a 1:1 ratio (red). **B.** Relative NMR signal intensity changes on FapB resonances due to the addition of equimolar unlabeled FapC at 1:1. The red curve on top indicates averages over 4 residues. **C.** Chemical shift perturbations (CSP) at FapB NMR resonances due to the addition of equimolar unlabeled FapC at 1:1. Nine residues indicated strong CSPs, larger than mean+2STD, and are indicated in green, whereas moderate CSPs are colored in yellow. **D.** The plot of CSPs (strong and medium) on the FapB sequence/secondary-structure elements from the AF3 model of the cryo-EM structure presented here. **E.** The plot of CSPs (strong and medium) on the FapB AF3 model, along with the AF3 model’s electrostatic surface representation.

To quantify interaction-induced perturbations, CSPs were calculated as Δδ = (Δδ = ((5*Δδ^1H^)^2^ + (Δδ^15N^)^2^)^1/2^) and categorized as negligible (<0.047 ppm), moderate (0.047–0.07 ppm), or strong (>0.07 ppm). Overlaying the HSQC spectra showed that most FapB peaks remained unchanged in both position and intensity (**Fig. 4A-C**). No new resonances or widespread line broadening were observed, indicating that FapB does not form a stable or highly populated monomer-monomer complex with FapC under these conditions. These data do not exclude weak, rapidly exchanging, or sparsely populated transient interactions below the sensitivity of the experiment.

Nevertheless, seven residues in FapB showed strong CSPs: F88, A101, V124, T141, G143, S144 and G178. The largest shifts were observed for V124 (0.124 ppm) and S144 (0.123 ppm). These small and localized CSPs do not indicate a large-scale FapC-induced conformational change, as we plotted the CSPs on the AF3 model and cryo-EM structure we present here (**Fig. 4E**). Importantly, the predicted β-strand regions of FapB remained largely unperturbed upon FapC addition, providing no evidence that monomeric FapC induces early folding or conformational priming of monomeric FapB (**Fig. 4D**).

### FapB and FapC exhibit distinct self- and cross-seeding behaviors

To evaluate how preformed fibril seeds influence amyloid nucleation and elongation, we conducted self-seeding (monomer + matched seed) and cross-seeding (monomer + nonmatched seed) experiments using 40 μM monomers with 5% or 20% sonicated fibril seeds (molar percentage) (**Fig. 5**). Aggregation kinetics were tracked via ThT fluorescence, and seed-only background signals were subtracted from all traces. Both FapB and FapC showed enhanced fibrillation upon seeding, but with different sensitivities and seeding preferences.

**Fig. 5:**
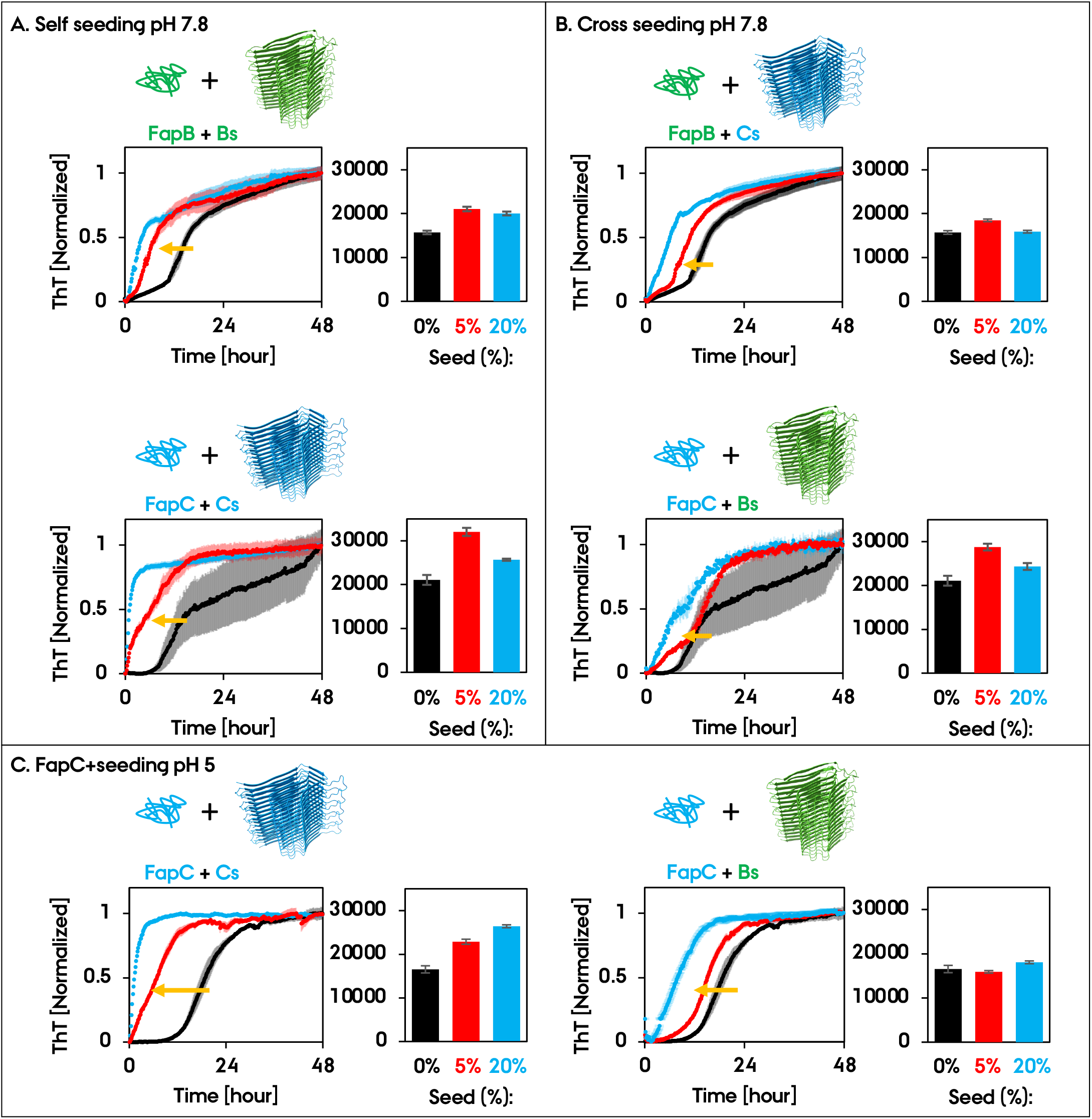
FapB is a nucleator for FapC & FapB, and FapC can successfully cross/self-seed. **A.** The ThT fibrillation assay results of self-seedings: FapB monomers (green IDP) with preformed FapB seeds (green filament) and FapC monomers (blue IDP) with preformed FapC seeds (blue filament) at pH 7.8. **B.** The ThT fibrillation assay results of cross-seedings, FapB monomers with preformed FapC seeds and FapC monomers with preformed FapB seeds at pH 7.8. **C.** The ThT fibrillation assay results of self-seedings, FapB monomers with preformed FapB seeds and FapC monomers with preformed FapC seeds at pH 5.0. The ThT assays are normalized to visualize the effect of seeding on the T_lag_ and fibrillation rate. The ThT_max_ changes are given at the side of each plot for no-seed (0%), as well as 5% and 20% seed ratios.

For FapB, both self- and cross-seeding accelerated fibrillation and increased ThT_max_, though self-seeding had the stronger effect (**Fig. 5A,B**). Unseeded FapB showed a T_lag_ of 1 h, T_50_ of 15 h, and ThT_max_ of ∼16,000. Self-seeding with 5% FapB fibrils shortened the T_lag_ to 0.5 h and T_50_ to 7.5 h and increased ThT_max_ to ∼21,000. At 20% seed, these parameters improved further, with a T_lag_ of 0.3 h, T_50_ of 5 h, and ThT_max_ of ∼20,000. SDS-PAGE confirmed greater monomer depletion 80-85% in seeded samples compared with ∼60% for unseeded controls (**Fig. SI10**).

Cross-seeding FapB monomers with FapC fibrils also accelerated fibrillation, albeit less efficiently than FapB self-seeding. Cross-seeding FapB monomers with 5% FapC seed shortened the FapB T_lag_ to 0.5 h and T_50_ to 11 h and increased ThT_max_ to ∼19,000. At 20% seed, these values improved further, with a T_lag_ of 0.3 h, T_50_ of 7.5 h, and ThT_max_ of ∼16,000. While cross-seeding with FapC clearly promoted FapB fibrillation, the effect was weaker than self-seeding at equivalent seed concentrations, indicating less efficient templating by FapC seeds compared with FapB seeds.

FapC displayed a distinct seeding response (**Fig. 5A,B**). Unseeded FapC showed a T_lag_ of 6 h, T_50_ of 18 h, and ThT_max_ of ∼21,000. Cross-seeding with FapB fibrils modestly improved kinetics. At 5% FapB seed, the T_lag_ decreased to 1 h and T_50_ to 14 h, while ThT_max_ increased to ∼29,000. At 20% FapB seed, these values improved further, with a T_lag_ of 1 h, T_50_ of 9 h, and ThT_max_ of ∼24,000. In contrast, FapC self-seeding had a much stronger effect. Addition of 5% FapC seed shortened the T_lag_ to 0.5 h and T_50_ to 6.5 h. At 20% FapC seed, the T_lag_ decreased to 0.3 h and T_50_ to 1 h. Interestingly, 5% FapC seeding gave the highest ThT_max_ of ∼32,000, exceeding the 20% self-seeded sample and all cross-seeded conditions. This stronger self-seeding response is consistent with the sequence alignment analysis, in which FapC L1 and FapC L3 aligned better than FapB L1 and FapC L3 **(Fig. SI4B).** SDS-PAGE showed >90% monomer depletion in all cases, indicating that seeding modulated kinetics but not the final extent of monomer conversion (**Fig. SI11**)

A similar seeding response was observed when FapC fibrillation was repeated at pH 5 (**Fig. 5C**). Unseeded FapC showed a T_lag_ of 7 h, a T_50_ of 18.5 h, and a ThT_max_ of ∼17,000. Cross-seeding with FapB fibrils accelerated fibrillation, with 5% FapB seed reducing the T_lag_ to 3 h and the T_50_ to 15 h, while the ThT_max_ remained similar at ∼16,000. Increasing the FapB seed to 20% further reduced the T_lag_ and T_50_ to 2 h and 8 h, respectively, with a modest increase in ThT_max_ to ∼18,000. FapC self-seeding produced a more pronounced acceleration, with 5% FapC seed reducing the T_lag_ to 0 h and the T_50_ to 7 h, accompanied by an increase in ThT_max_ to ∼23,000. At 20% FapC seed, the T_50_ was further reduced to 1.2 h, while the T_lag_ remained at 0 h and ThT_max_ increased to ∼26,000. Consistent with the results at pH 7.8, SDS-PAGE showed >95% monomer depletion across all conditions, indicating that seeding primarily accelerated fibrillation kinetics without substantially altering the extent of monomer conversion (**Fig. SI12**).

### FapB functional amyloid adopts a three-layer β-solenoid fibril structure by Cryo-EM

To determine the structural organization of FapB fibrils, we performed negative-stain EM and cryo-EM (**Fig. 6**, **Figs. SI7, SI13**). Negative-stain EM revealed relatively homogeneous FapB fibrils that were narrower than FapC fibrils, with diameters of approximately 7.2 and 8.2 nm, respectively (**Fig. SI7**). Cryo-EM 2D class averages showed well-defined single-protofilament fibrils with a regular axial repeat characteristic of an ordered amyloid architecture, corresponding to a strong layer line at 1/4.7 Å in the power spectrum (**Fig. SI7**). Compared with FapC, FapB exhibited a slightly narrower ordered core and a less prominent peripheral “fuzzy coat” (**Figs. SI7**), consistent with shorter flexible loop and terminal regions. This interpretation is also consistent with the AF3 model (**Figs. 1A, SI2, SI7**), which predicts a compact FapB cross-β-solenoid with shorter loops and relatively limited terminal extensions.

**Fig. 6:**
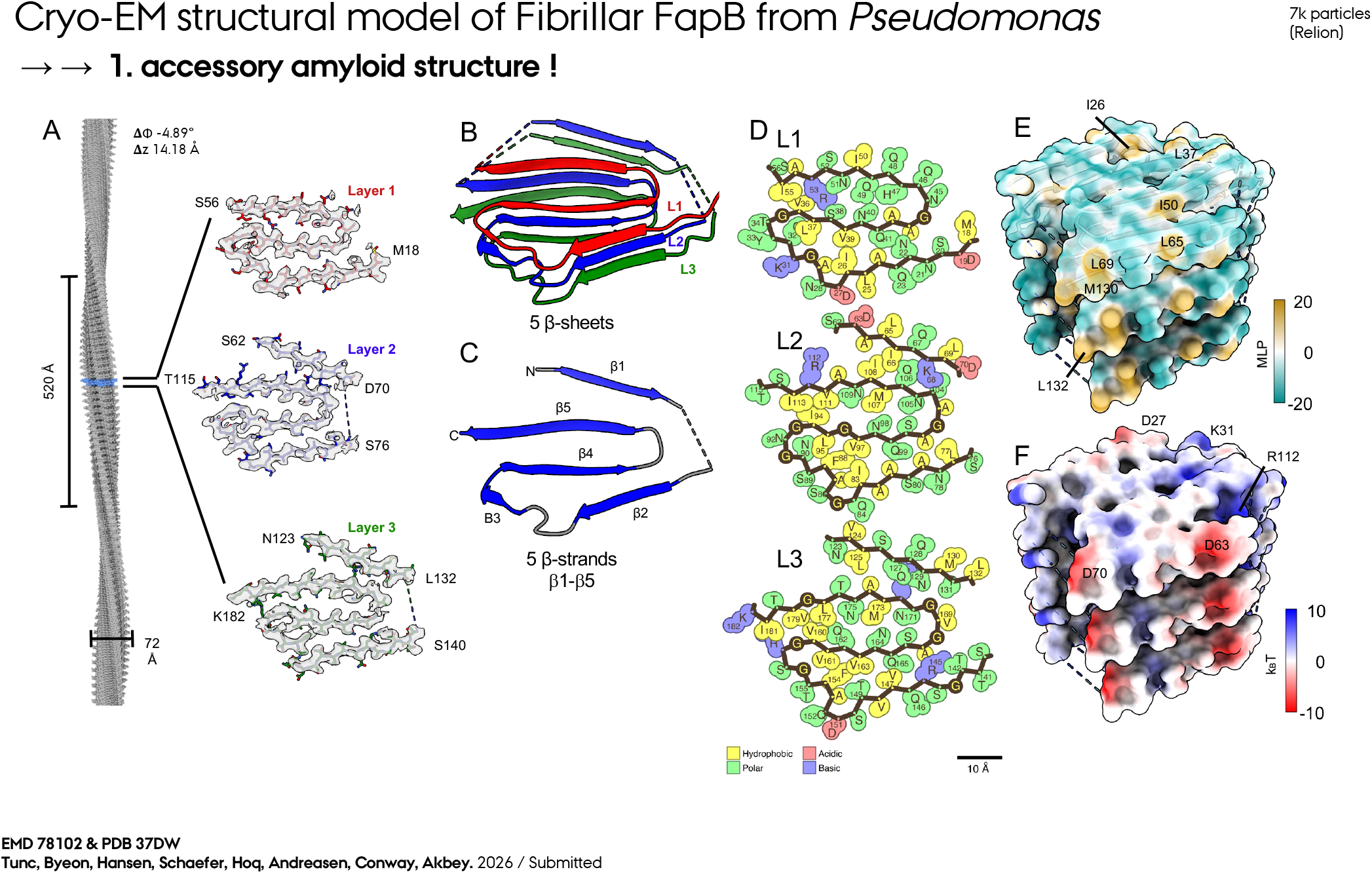
Cryo-EM reconstruction of FapB accessory functional amyloid. **A.** Symmetry-expanded helical model of FapB amyloids with a diameter of 72 Å, cross-over distance of 520 Å, helical rise (**Δ**z) of 14.18 Å, and helical twist (**Δ**ɸ) of −4.89°. Per-layer map-model fit shown in grey, with layer L1 in red, L2 in blue, and L3 in green. **B.** Overview of the FapB subunit with annotated layer structure. **C.** Consensus secondary structure annotation using layer L2, loops in grey, strands in blue for β1-β5 strands. **D.** Per-layer and per-residue property annotation from Amyloid Illustrator, indicating different amino acid types.(*49*) **E.** Molecular lipophilicity potential (MLP) colored surface with key lipophilic residues annotated. **F.** Surface electrostatics colored model with key polar residues annotated. Calculations were performed in ChimeraX using the APBS function.

To assess the robustness of the FapB reconstruction, we processed the same dataset using two parallel helical reconstruction workflows: our previously established cryoSPARC-based pipeline used for PSMα1 and FapC, (*15, 17, 47*) and an independent RELION5-based reconstruction strategy (**Fig. SI13**).(*48*) The cryoSPARC workflow yielded an initial FapB-UK4 reconstruction with a nominal overall resolution of ∼3.1 Å (**Fig. SI13**). Despite this higher nominal global resolution, local discontinuities in the backbone density and limited side-chain definition reduced its suitability for atomic interpretation. Our approach highlights previously identified limitations of classifying symmetry-related helical assemblies in cryo-SPARC, pointing to the need for further software optimization.(*47*) Therefore, independent helical reconstruction in RELION5 using a conventional amyloid helical reconstruction workflow (**Fig. SI13**),(*48*) produced a density map at an overall resolution of ∼3.4 Å with improved backbone continuity and side-chain definition. We therefore used the RELION reconstruction for subsequent structural interpretation and model analysis (**Fig. 6A**).

FapB-UK4 predominantly assembled into fibrils with a diameter of ∼70 Å and a helical crossover distance of ∼520 Å. The dominant morphology represented approximately ∼70% of the classified fibril segments, whereas an additional ∼30% displayed more highly twisted morphologies with shorter crossover distances (**Fig. SI13**). These minor classes did not yield reconstructions of sufficient quality for high-resolution structural determination, indicating limited structural polymorphism within the FapB fibril population. The dominant FapB fibril was refined using a helical rise of Δz = 14.18 Å and a twist of Δϕ = −4.89° per three-layer FapB subunit (**Fig. 6A**). The absolute handedness was not experimentally established and therefore remains formally ambiguous, although the reconstruction parameters are consistent with those of the previously determined left-handed FapC fibrils (**Fig. 6**).

Similar to FapC, the FapB fibril is organized as a three-layer β-solenoid. The ordered regions define three consecutive layers, L1 (M18–S56), L2 (S62–T115), and L3 (N123–K182) (**Fig. 6A,B**). Flexible segments connecting these layers are less well resolved than the β-sheet-rich core. Because helical averaging can obscure local differences among equivalent layers, the precise trajectories of some layer-crossing regions and unambiguous assignment of individual side chains remain less certain. The reconstruction nevertheless supports a continuous backbone throughout most of the β-solenoid core and exhibits an overall mean Q-score of 0.53 between FapB model and the helical reconstruction from RELION (**Fig. SI13**). Some connecting and peripheral regions remain less certain because of reduced local density, limiting unambiguous side-chain assignment in these segments.

Layers, L1-L3, are composed of five β-strands, and these form five β-sheets along the fiber z-axis. Notably, L1 lacks the β-strand corresponding to β1 in L2 and L3 at the sequence/structural-model level, yet the experimental density does not display a comparable discontinuity at this position. The β1-associated density in L2 and L3 is comparatively weak and likely conformationally heterogeneous. We therefore interpret the apparent continuity of this region cautiously rather than assigning it to a specific alternative strand register or averaging mechanism. This local variability contrasts with the substantially better resolved β-sheet-rich core. The C-terminal purification tag may also contribute partially to some of the additional density for L1 from an adjacent monomer.

Residue-level analysis of the FapB cryo-EM structure using Amyloid Illustrator revealed closely packed,(*49*) steric-zipper-like interfaces between five β-sheet surfaces despite being very close to each other at the sidechains of opposing layers/β-sheets, particularly involving β1-β5, β2-β4 and β4-β5, with stabilization dominated by hydrophobic side-chain contacts (**Fig. 6C,D**). Among the three layers, the central L2 layer contains the greatest density of hydrophobic residues, whereas L1 and L3 contain a larger proportion of polar residues. Molecular lipophilicity potential mapping therefore reveals a comparatively hydrophobic central region surrounded by more polar surfaces (**Fig. 6D,E**). Electrostatic surface analysis further shows that charged residues are enriched in solvent-exposed regions, particularly around the less well-resolved β1/2 segments, consistent with the increased conformational variability observed in these regions (**Fig. 6F**).

We next evaluated the energetic contributions of individual layers and residues using solvation-energy calculations implemented in Amyloid Illustrator (**Fig. 7A,E**). The central FapB layer L2 contains extended regions of favorable interactions and exhibits a calculated ΔG° of approximately −6.6 kcal mol⁻¹ layer⁻¹, compared with −0.9 and -4.1 kcal mol⁻¹ layer⁻¹ for L1 and L3, respectively (**Fig. SI7E, SI14**). Thus, stabilization is distributed unevenly across the three-layer repeat, with L2 making the strongest calculated energetic contribution. The central layer forms a hydrophobic core, whereas the flanking layers provide complementary packing interactions and solvent shielding. The calculated mean stability for full-length FapB was approximately −0.25 kcal mol⁻¹ residue⁻¹, comparable to values obtained using the same analysis for the two FapC structures, −0.18 kcal mol⁻¹ residue⁻¹ for PDB 9NQD and −0.20 kcal mol⁻¹ residue⁻¹ for PDB 9U4U. Similar values are observed for several other functional amyloid structures, including the FUS core (PDB 5W3N), hnRNPA2 LCD (PDB 6WQK), and HET-s prion (PDB 2RNM).(*49*) By comparison, selected pathological amyloid structures exhibit stronger calculated stabilization per residue, including an Aβ polymorph (∼-0.75 kcal mol⁻¹ residue⁻¹; PDB 5KK3) and a tau polymorph (∼-0.5 kcal mol⁻¹ residue⁻¹; PDB 6TJX).(*49*) These comparisons suggest that the extended FapB architecture can achieve substantial overall fibril stabilization through a large number of individually moderate interactions distributed throughout the β-solenoid.

**Fig. 7:**
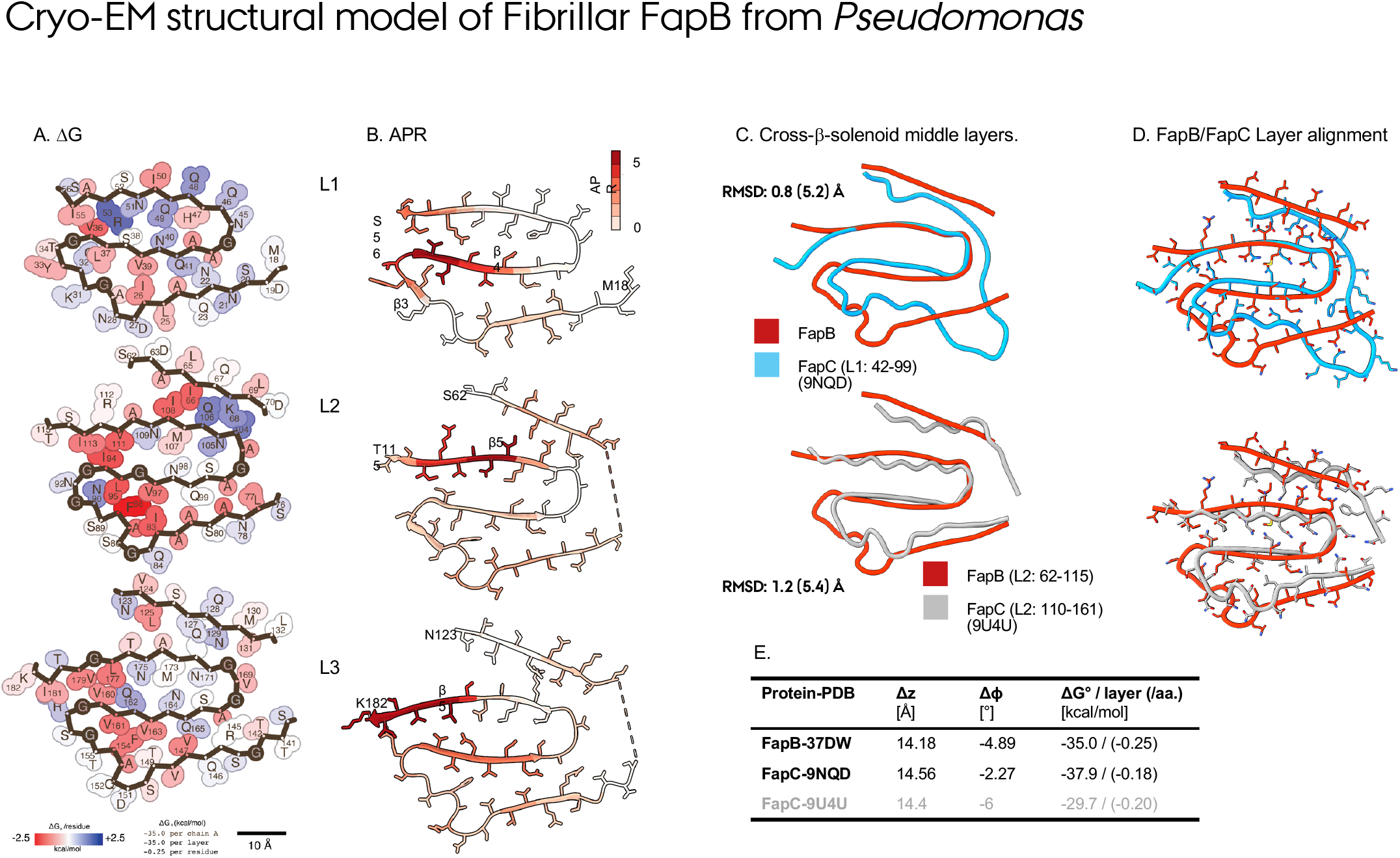
Properties of FapB accessory functional amyloid. **A.** Per-layer and per-residue amyloid solvation energies (**Δ**G°) from Amyloid Illustrator. Red: stabilizing; blue: destabilizing residue. **B.** Per-layer and per-residue calculation of amyloid-prone regions (APR) using a summed method score of AmylPred2.(*50*) **C.** Structural superposition of the cross-β-solenoid L2 (middle layer) of FapB and two published reconstructions of FapC (9NQD – 9U4U). The RMSD values represent pairwise structure comparisons of the aligned (all residues) RMSD values. The residue numbers and layer indices for the FapC layers from two different structures are given to be compared to the FapB L2 residues 62-115. **D.** Residue-level sidechain superpositions of the middle layer (L2) of FapB with two FapC structures (L2 for 9U4U and L1 for 9NQD, differing due to different Cryo-EM structures). **E.** Combined helical parameters and solvation-energy values per layer and per-residue for current FapB and two FapC structures.

To identify sequence regions with elevated intrinsic amyloid propensity, we calculated amyloid-prone regions (APRs) using AmylPred2.(*50*) Consensus scores from six sequence-based prediction methods were mapped onto the experimentally determined FapB structure (**Fig. 7B**). Predicted APRs occur in all three β-solenoid layers, including β2– β3 within L1, β4 within L2, and the C-terminal β4 region of L3. Thus, predicted amyloid propensity is distributed throughout the FapB sequence rather than concentrated within a single short segment, consistent with formation of an extended β-solenoid architecture.

Finally, we compared the FapB fold with experimentally determined FapC structures to assess structural compatibility between these two extracellular functional amyloids. Because the three layers contain related but nonidentical sequence repeats, we used the central FapB L2 sequence (SLNASAAIQGSAFSNGNGILGVNQSAGANNQMINAVRIST) as a representative layer and compared it with the corresponding structural regions of FapC, using L1 from PDB 9NQD and L2 from PDB 9U4U (**Fig. 7C**). Superposition with FapC PDB 9U4U yielded a Cα RMSD of approximately 1.3 Å within the aligned amyloid core of the two, whereas comparison with FapC PDB 9NQD yielded a Cα RMSD of approximately 0.8 Å. When the complete corresponding layers, including the more variable peripheral regions, were considered, the RMSDs increased to approximately 4.9 and 7.2 Å for 9U4U and 9NQD, respectively. Thus, FapB and FapC share a remarkably conserved β-solenoid core geometry, in particular at the β3-β5 core region (**Fig. 7D**), despite substantial divergence in the surrounding loops and peripheral regions (**Fig. 7C**,D). This conservation suggests that the underlying amyloid-core architecture can accommodate considerable sequence and peripheral structural variation while preserving a compatible fibril framework. Such structural compatibility may help explain the ability of FapB and FapC to participate in the same extracellular amyloid system and provides a plausible molecular basis for their observed cross-seeding behavior.

## DISCUSSION

Our findings reveal how a conserved amyloid architecture can support distinct and environmentally tunable assembly behaviors within the multicomponent *Pseudomonas* Fap system. The cryo-EM structure of *P. fluorescens* UK4 FapB establishes that this minor extracellular fibril component forms an independently stable three-layer cross-β-solenoid closely related to the major fibrillar protein FapC. Despite this structural conservation, FapB and FapC exhibit markedly different fibrillation kinetics, respond differently to pH and stoichiometry, and preferentially self-template while retaining reciprocal cross-seeding activity. Importantly, monomeric FapC does not induce large-scale conformational reorganization of intrinsically disordered FapB, indicating that molecular recognition between the two proteins becomes substantially more pronounced at higher-order stages of amyloid assembly. Together, these observations support a model in which FapB is an independently amyloidogenic but kinetically distinct component of the Fap system that can contribute to nucleation and templating in a condition-dependent manner rather than acting through a rigid, universal assembly hierarchy.

### A conserved β-solenoid core accommodates substantial structural divergence

At the molecular level, FapB adopts a compact three-layer β-solenoid that closely resembles the overall architecture of FapC despite their limited sequence identity. The experimentally determined FapB structure is in broad agreement with the AF3 prediction, with the most substantial deviations occurring in flexible loops, layer-connecting regions, and terminal segments (**Fig SI2**). FapB is narrower than FapC and exhibits a less prominent peripheral fuzzy region, consistent with its shorter loops and reduced terminal extensions. In addition, FapB L1 lacks the β-strand corresponding to β1 in L2 and L3, introducing a degree of asymmetry absent from the more regular FapC repeat architecture. Nevertheless, one dominant FapB fibril morphology yielded the high-resolution reconstruction, with only a limited population of more highly twisted polymorphs, indicating that FapB combines a well-defined principal fold with modest structural heterogeneity. We acknowledge that specific polymorph distributions may vary between current *in-vitro* fibrils and in native *Pseudomonas* biofilm. Nevertheless, we showed for FapC that the *in-vitro* and *ex-vivo* biofilm-extracted fibrils have similar dimensions.(*17*)

The structural comparison with FapC is particularly informative. Superposition of the aligned amyloid cores yielded Cα RMSDs of approximately 0.8-1.3 Å, whereas inclusion of the complete corresponding layers increased the RMSDs to approximately 4.9-7.3 Å (**Fig. 7C,D**). Thus, FapB and FapC preserve a highly similar amyloid-core geometry while tolerating considerably greater variation in peripheral and loop regions. This separation between a conserved structural core and divergent external regions provides a plausible framework for understanding how different Fap proteins can remain mutually compatible while retaining distinct kinetic and surface properties. The conserved core may preserve the geometry required for amyloid templating, whereas variation in solvent-exposed loops, termini, and electrostatic surfaces could tune partner recognition, fibril morphology, and assembly efficiency.

The energetic analysis further supports this view: FapB stabilization is distributed unevenly across the three-layer repeat, with the central L2 providing the strongest calculated energetic contribution and a comparatively hydrophobic interior, whereas the flanking layers provide complementary packing and more polar solvent-exposed surfaces. The calculated mean stabilization of FapB is similar to that of the experimentally determined FapC structures and several other functional amyloids, but weaker per residue than selected pathological amyloid structures.(*44*) These comparisons are consistent with an extended β-solenoid in which substantial overall fibril stability emerges from a large number of individually moderate contacts distributed across the full protein rather than from a single exceptionally stable short amyloid core like Aβ.

### Monomeric state, sample purity, and tag orientation shape FapB fibrillation

FapB is intrinsically amyloidogenic and assembles through a primary-nucleation-elongation-dominated pathway. A central finding of this study is that purely monomeric FapB is itself competent to form amyloid fibrils. SEC-purified FapB assembled robustly without detectable pre-existing oligomers. Our findings show that FapB fibrillation is strongly dependent on the starting sample state because Ni-NTA-purified samples containing HMW oligomers and minor LMW species fibrillated substantially more slowly and less efficiently than isolated monomer. This emphasizes that amyloid behavior cannot be interpreted independently of protein preparation. Similar sensitivity to oligomeric heterogeneity was previously observed for FapC, although individual oligomeric populations affected FapC differently.(*40*) Thus, high apparent purity by SDS-PAGE is insufficient to define an amyloid-forming preparation mechanistically, and even minor heterogeneous species can markedly alter the observed kinetic behavior.

Affinity-tag placement provided an additional example of this sensitivity. C-terminally tagged UK4 FapB fibrillated efficiently, whereas N-terminal tagging substantially delayed assembly. This behavior contrasts with that reported for PAO1 FapB, where an N-terminally tagged construct was favorable.(*42*) Although the molecular origin of this difference remains unresolved, it demonstrates that affinity tags are not necessarily mechanistically neutral in Fap proteins and may perturb local charge, steric accessibility, terminal conformational preferences, or interactions involved in early assembly. Such construct-dependent effects should be considered carefully when comparing Fap fibrillation across strains or studies.

### Distinct kinetics and reciprocal seeding define a tunable nucleator function

When strictly monomeric FapB and FapC were compared under identical conditions, their kinetics separated two distinct aspects of amyloid formation. FapB entered detectable assembly earlier, exhibiting a shorter apparent lag phase, but subsequently reached half-maximal fluorescence more slowly and retained a larger soluble monomer fraction. FapC, in contrast, exhibited a somewhat longer initial lag phase but underwent more rapid fibril accumulation and near-complete monomer conversion once assembly began. These results suggest that the energetic requirements for entering an assembly-competent state and for sustaining efficient fibril growth are not equivalent. FapB appears able to cross the initial nucleation barrier relatively readily under these conditions with short T_lag_, whereas FapC is more efficient in later sustained monomer incorporation and bulk scaffold formation with larger T_max_, defining different behavior and possibly different roles of assembly to individual Fap proteins.

The low-level seeding experiments provide additional mechanistic insight. Addition of 1-100 nM preformed FapB or FapC fibrils to FapC monomer did not measurably shorten the lag phase or accelerate fibrillation. Consistent with this observation, global analysis of the concentration-dependent FapB and FapC kinetics using AmyloFit was adequately described by a primary nucleation-elongation model, whereas inclusion of more complex secondary-nucleation pathways did not improve the quality of the fits. These results argue against a dominant contribution of fibril-surface-catalyzed secondary nucleation under the conditions examined. This behavior differs from many pathological amyloid systems in which secondary nucleation can provide a powerful autocatalytic route for fibril proliferation.(*27*) Importantly, however, the absence of a strong response to very low seed concentrations does not imply that fibrils are incapable of templating assembly. Rather, the higher-seed experiments demonstrate that FapB and FapC fibrils become effective templates when sufficiently abundant, distinguishing fibril-end primary-nucleation-elongation from a secondary-nucleation-dominated amplification mechanism.

### Under physiological conditions, FapB-FapC recognition is strongly dependent on molecular state

Solution NMR provides an important constraint on how FapB and FapC interact before fibrillation as IDP monomers. Addition of equimolar monomeric FapC caused no widespread chemical-shift perturbations, line broadening, or appearance of new FapB resonances. Only a small number of residues exhibited localized CSPs, indicating that FapC does not induce global folding or large-scale conformational rearrangement of the IDP FapB monomer. These data further argue against formation of a stable or highly populated FapB-FapC monomeric complex under the conditions examined, although weak, transient, or sparsely populated interactions cannot be excluded.

This weak interaction at the monomeric level contrasts with the clear templating behavior observed once fibrils are present. At higher seed fractions, both FapB and FapC strongly preferred homologous seeds but also exhibited reciprocal cross-seeding. FapB fibrils accelerated FapC assembly, and FapC fibrils accelerated FapB, although self-seeding remained more efficient in both directions. These results indicate that partner recognition within the Fap system is strongly dependent on conformational state: the soluble proteins remain largely independent, whereas the ordered fibrillar state generates interfaces capable of cross-templating structurally related partners.

The cryo-EM structures provide a plausible molecular explanation for this behavior (**Fig. 1A,7C,D**). The highly conserved β-solenoid core geometry of FapB and FapC could lower the structural mismatch required for heterologous templating, whereas differences in loops, termini, surface charge, and local β-strand organization may account for the preference for homologous seeds (**Fig. 7C,D**). In this view, self-seeding benefits from optimal surface complementarity, while cross-seeding remains possible because both proteins share a closely related amyloid-core framework, with a smaller RMSD between structures (**Fig. 7D**). Direct visualization of seed-monomer interfaces or structurally resolved heteromeric fibrils will ultimately be required to determine whether cross-seeding involves templating at fibril ends, interactions with lateral fibril surfaces, transient heterotypic nuclei, or combinations of these mechanisms.

Together, the NMR and seeding experiments therefore support a two-stage recognition model in which FapB and FapC remain predominantly disordered and conformationally independent as soluble monomers but become functionally coupled after amyloid surfaces are generated. Such state-dependent recognition provides a potential mechanism for limiting premature association while preserving the ability of extracellular Fap components to interact efficiently during matrix assembly.

### pH and stoichiometry tune the FapB-FapC assembly landscape

Environmental conditions strongly modulated the behavior of both proteins. At pH 5.0, FapB exhibited a substantially longer lag phase than at pH 7.8, yet its final monomer conversion increased markedly. FapC similarly fibrillated more slowly and produced a lower ThT response under acidic conditions while still undergoing near-complete monomer depletion. These observations further illustrate that lag time, ThT intensity, and final monomer conversion report different aspects of the assembly process and should not be interpreted interchangeably. Acidic conditions can therefore delay the onset of detectable fibrillation while simultaneously favoring more complete incorporation of FapB into insoluble assemblies.

The relative abundance of FapB and FapC imposed an additional level of regulation. Near-neutral conditions permitted efficient assembly over a comparatively broad range of FapB:FapC ratios, particularly when FapC was present in excess. At acidic pH, this permissive regime narrowed considerably, with strongly asymmetric mixtures showing reduced total incorporation of both components and the 1:1 condition supporting the most effective combined conversion. These findings indicate that pH modifies the range of stoichiometries over which the two proteins can assemble efficiently, potentially through changes in protonation, electrostatic complementarity, conformational populations, or fibril-surface interactions. Our mixed-monomer experiments nevertheless do not establish that FapB and FapC form heteromeric fibrils. The nonadditive kinetic behavior and simultaneous depletion of both proteins demonstrate that the assembly pathways influence one another, but the endpoint could contain heteromeric fibrils, separate populations of FapB and FapC fibrils, or a mixture of both. Direct structural and compositional characterization of fibrils formed from mixed monomers will therefore be required to determine whether FapB is incorporated into FapC-rich fibrils, functions primarily as a transient nucleator, or populates a separate but kinetically coupled fibril population.

The behavior observed when FapC is present in excess is particularly relevant to the known composition of native Fap fibrils. Previous biochemical analyses of extracellular fibrils from *Pseudomonas* showed that FapC constitutes the major structural component, whereas FapB and FapE are present at substantially lower abundance.(*13,14*) Moreover, disruption of FapA dramatically changes this composition: highly stable amyloid aggregates are still formed but become dominated by FapB with only a minor FapC fraction in the extracellular matrix, indicating that the relative incorporation of the two amyloidogenic proteins is actively regulated rather than fixed.(*14*) Conversely, FapE is present only at low abundance within mature fibrils yet is required for efficient extracellular amyloid production, supporting an important role in Fap biogenesis or secretion rather than simply serving as a major structural subunit.

### A flexible rather than universal hierarchy may govern Fap assembly

Our observations also highlight potential variation in Fap assembly behavior between *Pseudomonas* strains. Previous work with PAO1 Fap proteins reported a more asymmetric relationship in which FapC promoted FapB fibrillation whereas FapB had comparatively little influence on FapC.(*42*) In contrast, UK4 FapB and FapC display reciprocal cross-seeding, although homologous templating remains preferred. Sequence variation among FapB homologs is concentrated largely within loop and terminal regions rather than the conserved β-solenoid core, providing a plausible route by which different strains could preserve the overall amyloid architecture while altering fibril-surface chemistry and partner compatibility.

However, strain-specific sequence differences are unlikely to be the only possible explanation. The strong effects of purification state, affinity-tag position, pH, protein concentration, and stoichiometry observed here demonstrate that Fap kinetics are highly context-dependent. Differences between UK4 and PAO1 could therefore arise from sequence variation, experimental conditions, construct design, or combinations of these factors. Rather than establishing a single strain-specific mechanism, the current data argue against a rigid universal Fap nucleation hierarchy and instead support a more flexible assembly landscape in which conserved structural compatibility is modulated by both protein sequence and environment.

Multicomponent AF3 modeling provides a testable structural hypothesis for how such an assembly system might be organized. The models repeatedly position FapE in contact with FapB/FapC and place FapB between FapE and FapC structures, raising the possibility that FapB could participate in coupling early FapE-dependent biogenesis events to subsequent FapC-rich fibril growth. This arrangement is intriguing considering the biochemical observation that FapE is a low-abundance but essential component of Fap amyloid production and that FapB can become a dominant fibril component when FapA regulation is perturbed.(*19*) Nevertheless, neither the AF3 models nor the present *in vitro* experiments establish the temporal sequence, stoichiometry, or physical composition of such a FapE-FapB-FapC assembly intermediate. This discussion should therefore be viewed as an assembly hypothesis for future testing rather than as a resolved assembly pathway.

Overall, our results establish FapB as an independently amyloidogenic, structurally defined component of the Fap system whose behavior cannot be reduced to that of a passive minor fibril protein. FapB and FapC preserve a highly conserved β-solenoid core yet differ substantially in their peripheral architecture, fibrillation kinetics, conversion efficiency, and environmental sensitivity. Their monomers remain largely conformationally independent, whereas their fibrillar states are mutually compatible and capable of reciprocal cross-templating. At the same time, the absence of strong acceleration at very low seed concentrations and the kinetic modeling support an assembly mechanism dominated by primary nucleation and elongation rather than extensive secondary nucleation under the conditions examined. Together, these findings support a model in which conserved amyloid-core geometry provides structural compatibility, while sequence variation, fibril-surface properties, stoichiometry, and extracellular conditions tune the relative contributions of FapB and FapC to matrix assembly. This combination of structural conservation and kinetic flexibility provides a molecular framework for understanding how multicomponent functional amyloids can be assembled and regulated within the *Pseudomonas* biofilm extracellular matrix.

### CONCLUSIONS

FapB from *P. fluorescens* UK4 forms a compact three-layer cross-β-solenoid that shares a highly conserved amyloid-core architecture with FapC while retaining distinct peripheral structural features. Despite this structural similarity, FapB and FapC exhibit different fibrillation kinetics, environmental sensitivities, and templating preferences. FapB is intrinsically amyloidogenic and can contribute to nucleation and templating in a condition-dependent manner, whereas FapC undergoes more efficient monomer conversion and is consistent with its role as the principal fibrillar scaffold. Low-level seeding and kinetic modeling further support an assembly mechanism dominated by primary nucleation and elongation under the conditions examined rather than by extensive fibril-surface-catalyzed secondary nucleation. FapB and FapC remain largely conformationally independent as soluble monomers but become functionally coupled in the fibrillar state, where both proteins preferentially self-template yet retain reciprocal cross-seeding activity. This state-dependent recognition provides a mechanistic framework in which a conserved β-solenoid core permits heterologous templating, while differences in loops, terminal regions, surface chemistry, stoichiometry, and pH tune assembly efficiency and partner compatibility. The contrasting behavior reported for PAO1 further suggests that Fap assembly does not follow a single rigid nucleation hierarchy and may instead be modulated by both strain-specific sequence variation and experimental or environmental context. Multicomponent AF3 models predicting a FapE-FapB-FapC organization provide a testable hypothesis in which FapB could link early FapE-dependent biogenesis events to subsequent FapC-rich fibril growth; however, the temporal order, stoichiometry, and coexistence of these proteins within individual fibrils remain to be established. Overall, our findings define FapB as an independently amyloidogenic and structurally conserved, yet kinetically distinct, component of the Fap system and illustrate how multicomponent functional amyloids can preserve a common structural framework while tuning molecular recognition and assembly in response to biological and environmental context.

## METHODS

### FapB and FapC Constructs

For the C-terminally His-tagged FapB construct, residues 19-182 were cloned into the pET28a vector system. This construct introduced one additional amino acid (M) at the N-terminus and nine additional amino acids (WLEHHHHHH) at the C-terminus. For the N-terminally His-tagged FapB construct, residues 19-182 were cloned into a modified pET28a vector designed to improve protein expression. In this construct, the thrombin cleavage site was replaced with a TEV protease cleavage site. The T7 promoter and translation initiation region were further modified using the T7p^CONS^ and TIR-2 sequences, respectively, as described previously to improve protein expression.(*51*) These modifications were introduced to enhance protein expression. The resulting construct contained 13 additional amino acids (MQLPWHHHHHHGS) at the C-terminus and no additional amino acids at the N-terminus. N-terminally and C-terminally His-tagged FapB has calculated molecular weights of 18.4 kDa and 18.2 kDa. The preparation of FapC 25-250 (FL), comprising residues 25-250 and lacking the signal sequence (residues 1-24), has been described previously in detail.(*40*)

### Protein Expression and Purification

Recombinant protein constructs were expressed in *Escherichia coli* BL21(DE3) cells using plasmids encoding residues 19-182 of FapB. The N-terminal signal peptides (residues 1-18 for FapB and 1-24 for FapC) were omitted from all constructs because they are cleaved during secretion and are not incorporated into functional fibrils *in-vivo*. ^15^N labeled FapB was used throughout the study, except for the experiments corresponding to Fig. SI6 and SI7, which used unlabeled protein. FapC used in all experiments is unlabeled.

For NMR-related experiments requiring isotopic labeling, cells were cultured in minimal medium containing ^15^N-ammonium chloride (1 g L⁻¹) as the sole nitrogen source, whereas unlabeled protein was produced using LB medium. Glycerol stocks were first used to inoculate overnight LB cultures containing the appropriate antibiotic, which were incubated at 37 °C with shaking. The resulting cultures were harvested and resuspended in fresh growth medium, followed by incubation at 37 °C until an optical density at 600 nm (OD600) of 1.0-1.5 was reached. Protein expression was initiated by adding isopropyl β-D-1-thiogalactopyranoside (IPTG) to a final concentration of 1 mM, after which cultures were incubated for an additional 1-2 h at 37 °C. Cells were then harvested by centrifugation at 6000 × g for 20 min at 4 °C and resuspended in denaturing lysis buffer containing 50 mM Tris (pH 8.0) and 8 M guanidinium chloride. Cell lysis was performed by sonication, and insoluble material was removed by centrifugation at 6000 × g for 20 min at 4 °C, resulting in a clarified soluble lysate.

Proteins were purified under denaturing conditions by Ni-NTA affinity chromatography using HisPur Ni-NTA resin (Thermo Fisher Scientific). Clarified lysates were incubated with the Ni-NTA resin overnight at 4 °C with gentle mixing to facilitate protein binding. The resin was subsequently collected by centrifugation, and the unbound material was removed. Extensive washing with lysis buffer was then performed to eliminate nonspecifically bound proteins. Bound FapB was eluted stepwise with lysis buffer containing increasing concentrations of imidazole (30, 60, 120, 300, and 500 mM). Elution fractions were collected individually and either analyzed separately or pooled as specified for subsequent experiments. The resulting protein samples were concentrated using centrifugal filter units with a 3 kDa molecular weight cutoff. Any subsequent size-exclusion chromatography used to separate monomeric from non-monomeric species is described in the corresponding sections below.

### Size-Exclusion Chromatography and Separation of Oligomeric States

Size-exclusion chromatography (SEC) was performed to separate monomeric proteins from oligomeric and truncated species before fibrillation assays. Following Ni-NTA affinity purification and concentration, protein samples were applied to a pre-equilibrated HiLoad 16/600 Superdex 200 column (Cytiva Life Sciences). Chromatographic separation was carried out under denaturing conditions using a SEC buffer consisting of 50 mM Tris (pH 8.0) and 8 M guanidinium hydrochloride to maintain the protein in its denatured state. Samples were loaded at concentrations optimized to preserve resolution between monomeric and non-monomeric species while minimizing column overloading. Protein elution was monitored by UV absorbance at 280 nm, and fractions were collected throughout the complete elution profile. Monomeric fractions were identified and isolated based on SDS-PAGE analysis.

### Mass Spectrometry (MS) data acquisition and processing

ESI LC-MS measurements were conducted using 1 μM protein samples on a Bruker Q-TOF instrument equipped with a reverse-phase AdvanceBio peptide guard column (Agilent Technologies). Mobile phases A and B consisted of 5% and 80% acetonitrile, respectively, with 0.01% formic acid. LC-MS spectra were processed using Bruker Compass Software, and maximum-entropy-based deconvolution was applied to the MS data to determine the M+ ion mass for each sample. Prior to each measurement, the instrument was calibrated using ESI Low Tuning Mix I (Agilent Technologies) to achieve a mass difference of 1.0 ppm.

### Aggregation Kinetics Assays

Protein samples stored under denaturing conditions (8 M guanidinium hydrochloride) were desalted into fibrillation buffer (20 mM sodium phosphate, pH 7.8, 0.02% sodium azide) for the experiments corresponding to Fig. 2, 5A-B, and SI6A, B. For experiments corresponding to Fig. 3 and 5C, samples were instead desalted into universal buffer as described previously (20 mM Tris-HCl, 20 mM Bis-Tris, and 20 mM sodium acetate, pH 5 and pH 7.8, 0.02% sodium azide).(*52*) Desalting was performed using PD MiniTrap G-10 desalting columns (Cytiva). Protein concentrations were determined by measuring UV absorbance at 280 nm using a NanoDrop spectrophotometer (Thermo Scientific), with the following extinction coefficients: 6.99 mM⁻¹ cm⁻¹ for both FapB N-term and FapB C-term, 10.1 mM⁻¹ cm⁻¹ for FapC. Unless otherwise indicated, samples were prepared at a final protein concentration of 40 μM in fibrillation buffer, with ThT added to a final concentration of 200 μM. A volume of 50 μL from each sample was transferred to black, clear-bottom 96-well plates (Corning 3881, Sigma-Aldrich). ThT fluorescence was measured using a Tecan Spark plate reader with excitation and emission wavelengths of 442 nm and 482 nm, respectively. Fluorescence measurements were collected at 10-min intervals for at least 48 h at 37 °C without shaking. Unless otherwise stated, all kinetic experiments were performed in triplicate. Background fluorescence was corrected by subtracting the signal from buffer containing 200 μM ThT.

For seeding experiments, previously fibrillated samples at concentrations of 40 µM or higher were used and diluted to a final concentration of 40 µM after fibrillation. Seed formation was performed by sonication of these fibril samples at 10% power for 30 s, repeated twice with a 10 s interval between sonication cycles. The resulting seeds were then added to ThT assay samples at the specified molar-to-molar ratios. For ThT analysis of the seeding experiments, both the seed background signal and the ThT background signal were subtracted from the measured fluorescence signals.

### SDS-PAGE Analysis and Monomer Quantification

Fibrillation was evaluated by SDS-PAGE using samples collected immediately before (PRE) and after (POST) ThT incubation. For the PRE samples, aliquots were combined with SDS loading buffer, denatured at 50 °C, and subsequently stored at 4 °C. POST samples were treated using the same procedure after fibrillation. PRE and POST samples were then analyzed under non-reducing conditions using 12% SDS-PAGE gels. The intensity of the monomer bands was quantified using Image Lab Software for Mac Version 6.1 (Bio-Rad) and reported as a percentage of the initial monomer intensity in the corresponding PRE lanes. When PRE and POST samples were analyzed on separate gels, the gels were normalized relative to one another using a ladder band.

### Solution-State NMR Spectroscopy

Uniformly ^15^N-labeled full-length FapB (residues 19-182, lacking the signal peptide and containing a C-terminal His-tag) and unlabeled FapC (residues 25-250, lacking the signal peptide and containing a C-terminal His-tag) were used for solution NMR experiments. NMR data were collected on a Bruker Avance III 600 MHz spectrometer fitted with a 5-mm triple-resonance TCI cryogenic probe. Protein samples were prepared in 20 mM sodium phosphate buffer (pH 7.8) supplemented with 0.02% sodium azide, 1 mM deuterated dextran sulfate sodium, and 10% (v/v) D₂O. Samples were prepared immediately before NMR data collection and loaded into 5-mm NMR tubes at a final volume of 420 μL. Spectral processing was carried out using TopSpin 3 (Bruker BioSpin), and peak assignments were performed manually in CCPNmr Analysis 3. Proton chemical shifts were referenced against DSS at (0 ppm), whereas ^15^N chemical shifts were indirectly referenced using the ^1^H frequency in accordance with established conventions.

### Negative-Stain Electron Microscopy

Negative-stain EM imaging was used to assess fibril morphology and abundance. Copper grids (400-mesh) coated with a continuous carbon film were glow-discharged with 25 mA current for 90 s prior to sample application. Approximately 3 μl of protein sample was applied to each grid and incubated for 10 s before excess liquid was removed by side blotting with filter paper. Grids were stained with 2% (w/v) uranyl acetate for 10 s, followed by side blotting and air drying. Micrographs were collected on a Tecnai TF20 TEM (Thermo Fisher Scientific) operated at 200 kV and equipped with a TVIPS XF416 CMOS camera. Images shown were not subjected to further image processing.

### Preparation, data collection and processing for cryo-EM

80 µM FapB was diluted 10-fold in Milli-Q water and used for cryo-EM. Quantifoil R 1.2/1.3 grids were glow-discharged under vacuum for 60 s at 25 mA in a Pelco easiGlow 91000 glow discharge cleaning system (Ted Pella). A volume of 3 µL of sample was applied to Quantifoil R 1.2/1.3 grids. The grids were incubated before being vitrified using a Vitrobot Mark IV. AutoGrids were assembled and immediately screened. Cryo-EM datasets were collected on a Titan Krios (Thermo Fisher) operating at 300 keV. Movies were recorded using a Falcon 4i direct electron detector (Thermo Fisher) at a nominal magnification of 165,000, corresponding to a pixel size of 0.74 Å. Movies were saved in the electron-event representation (EER) format and recorded at a total electron exposure of 50 e^−^ per Å^2^. All datasets were collected automatically using EPU (v.X, Thermo Fisher) with a defocus range of −0.5 to -2.5 μm. EPU’s fast exposure navigation was used to collect data at an 8 µm image shift. All datasets were processed using cryoSPARC (v.4.7),(*53*) and RELION5(*54*). The processing workflow is shown in supplementary **Figure S13**. A total of 17,272 movies were collected and dose-fractioned into 18 fractions (no upsampling), motion-corrected using patch-based motion correction, followed by patch-based CTF estimation. Micrographs with CTF fits above 4 Å and astigmatism above 600 Å were discarded. The remaining 13,897 accepted micrographs were used for further processing. A full set of non-default parameters is given in the processing workflow (supplementary **Fig. S13B**).

Filaments were traced in template-free mode using a 50-120 Å diameter range and 0.1 diameter separation. Fibrils with curvature above 0.0015 and sinuosity above 1.1 were rejected. The remaining segments were extracted in a 1024 px box, binned to 100 px. 3.4 million segments were subjected to 2D classification with 100 classes. The main polymorph, representing 1.2 million segments, was selected, and a crossover distance (CO) was approximated using 2D class averages in ImageJ.(*55*) Helix refinement without imposed symmetry and a cylindrical starting volume of 80 Å diameter was used. Using a minimum of 10 segments per micrograph, exposures were curated again. Picks were re-extracted using a 360 px box, binned 3-fold, with a Nyquist frequency of 4.3 Å. Following 2D classification, class averages with visible cross-beta stack separation were selected. The resulting 408 thousand segments were further refined, assuming a single amyloid-layer assembly (rise: 4.85 Å and 1.46° twist). Segments were re-extracted without binning and refined with the final three-layer amyloid symmetry, assuming left-handedness (rise: 14.35 Å, twist: -5.48 °). Alignment-free 3D classification in PCA mode resulted in a well-defined class with 122 thousand segments, which were subjected to another helical refinement. To optimize helical parameters, segments were converted to a .star file using pyem, (https://doi.org/10.5281/zenodo.3576630) and used for 3D classification in RELION5,(*54*) without alignment and with helical search enabled. 87 thousand segments were imported back into cryoSPARC and refined with a rise of 14.25 Å and twist of -4.75°. CTF refinement was performed, followed by a final helical refinement, resulting in an optimized rise of 14.25 Å and twist of -4.6 °. Local motion was corrected with reference-based motion correction, followed by another round of alignment-free 3D classification and a final helical refinement using a rise of 14.22 Å and twist of -4.94°. The overall Gold-Standard resolution at FSC 0.134 was reported as 3.5 Å and a B-factor of -11 Å^2^.

Here are the details of the independent processing in RELION5. Movies were gain-corrected, motion-corrected and dose-weighted using MotionCor2 (ref). Contrast transfer function (CTF) parameters were estimated from aligned, non-dose-weighted micrographs using CTFFIND-4.1 (ref). Filaments were automatically picked using Topaz (ref), and helical segments were extracted in 380-pixel boxes. De novo initial models were generated from selected 2D class averages spanning a complete helical crossover using *relion_helix_inimodel2d* in RELION 5. Initial references were low-pass filtered to 10 Å before 3D classification. Successive rounds of 3D classification were used to select a structurally homogeneous subset of segments, followed by 3D auto-refinement. Helical rise and twist were optimized during selected refinement stages. Bayesian polishing, CTF refinement and additional classification and refinement were subsequently performed to improve the reconstruction and remove suboptimal segments. The final refinement used ∼8k segments with a fixed helical rise of 14.18 Å and twist of -4.89°. Soft masks encompassing the central 10-30% of the box along the helical axis were used for post-processing. The final map was sharpened using standard RELION post-processing procedures, yielding a reported masked resolution of 3.4 Å.

### Model building, refinement, and validation

To aid model building, the final map was modified using EMReady2,(*56*) which improved interpretability during model building. The initial atomic models were generated with AlphaFold using modified UniProt P0DXF4 M-19-182-WLEHHHHHH, which was fitted into the densities as rigid bodies within ChimeraX and manually corrected using Coot (v.0.9.8.95),(*57*) and ISOLDE and iteratively refined using *phenix.real_space_refine* within Phenix (v.1.19).(*58*) Validation reports were automatically generated using MolProbity within Phenix.(*59*) Q-scores were calculated within ChimeraX (v.1.12).(*60*) The full validation report is provided in **Table S1**.

### Calculation of amyloid-prone regions

AmylPred2 was used to assess the amyloid-prone regions (APRs) from the FapB-UK4 sequence.(*50*) AmylPred2 failed to access WALTZ- and AggreScan, which were calculated separately. AggreScan-values were binarized to 1 for a4v values above −0.02. The AmyloidMutants webserver was not reachable. The summed score rather than consensus APR was calculated using the predicted APRs from WALTZ, AggreScan, Amyloidogenic Pattern, Average Packing Density, Beta-strand contiguity, Hexapeptide Conf. Energy, NetCSSP, Pafig, SecStr and TANGO. The resulting summed scores were utilized to populate the residue-based B-factor column of the protein structure file for visualization with ChimeraX v.1.10.(*61*)

### Solvation energy calculation

Per-residue and per-layer solvation energies (ΔG°) were calculated using the Amyloid Illustrator software and plotted by separating the three-layer structure of FapB for visualization.(*49*) A three-peptide-model was used as input to correctly calculate the shielded central amyloid peptide.

### Data availability

The cryo-EM map of FapB has been deposited in the Electron Microscopy Data Bank (EMDB) under accession number **EMD-78102**. The refined atomic model has been deposited in the Protein Data Bank (PDB) under accession number **37DW**.

## Acknowledgements

UA acknowledge support from the Department of Structural Biology, University of Pittsburgh School of Medicine (UPSOM) for access to the high-field NMR and EM facilities. UA acknowledges the start-up funds by the UPSOM and a Competitive Medical Research Fund from the UPMC Health System. JHS acknowledges funding for this research from the German Research Council project number 556478029. The Pittsburgh Center for Cryo-EM (RRID:SCR_025216) used for data collection in this project was supported, in part, by the University of Pittsburgh, the School of Medicine, the Department of Structural Biology, and the National Institutes of Health through grants S10OD019995 (JFC) and S10OD025009 (JFC). The content is solely the responsibility of the authors and does not necessarily represent the official views of the National Institutes of Health. MA acknowledges funding from Aarhus University Research Foundation NOVA (AUFF-E-2023-9-18).

## Author contributions

Writing—original draft: A.T., C.H.B., J.H.S., and Ü.A. Conceptualization: Ü.A. Investigation: A.T., C.H.B., J.H.S., K.H.H., M.A., M.R.H., and Ü.A. Writing—review and editing: A.T., C.H.B., J.H.S., K.H.H., M.A., M.R.H., J.F.C and Ü.A. Methodology: A.T., C.H.B., J.H.S., K.H.H., M.A., M.R.H., J.F.C and Ü.A. Resources: Ü.A. Funding acquisition: Ü.A. Data curation: A.T., C.H.B., J.H.S., K.H.H., M.A., M.R.H., J.F.C and Ü.A. Validation: A.T., C.H.B., J.H.S., K.H.H., M.A., M.R.H., J.F.C. and Ü.A. Supervision: Ü.A. Formal analysis: A.T., C.H.B., J.H.S., K.H.H., M.A., M.R.H., J.F.C. and Ü.A. Software: J.H.S., M.R.H., and Ü.A. Project administration: D.A.G., J.H.S., Ü.A. Visualization: A.T., C.H.B., J.H.S., K.H.H., M.R.H., and Ü.A.

## Competing interests

The authors declare that they have no competing interests.

**Fig. SI1:**
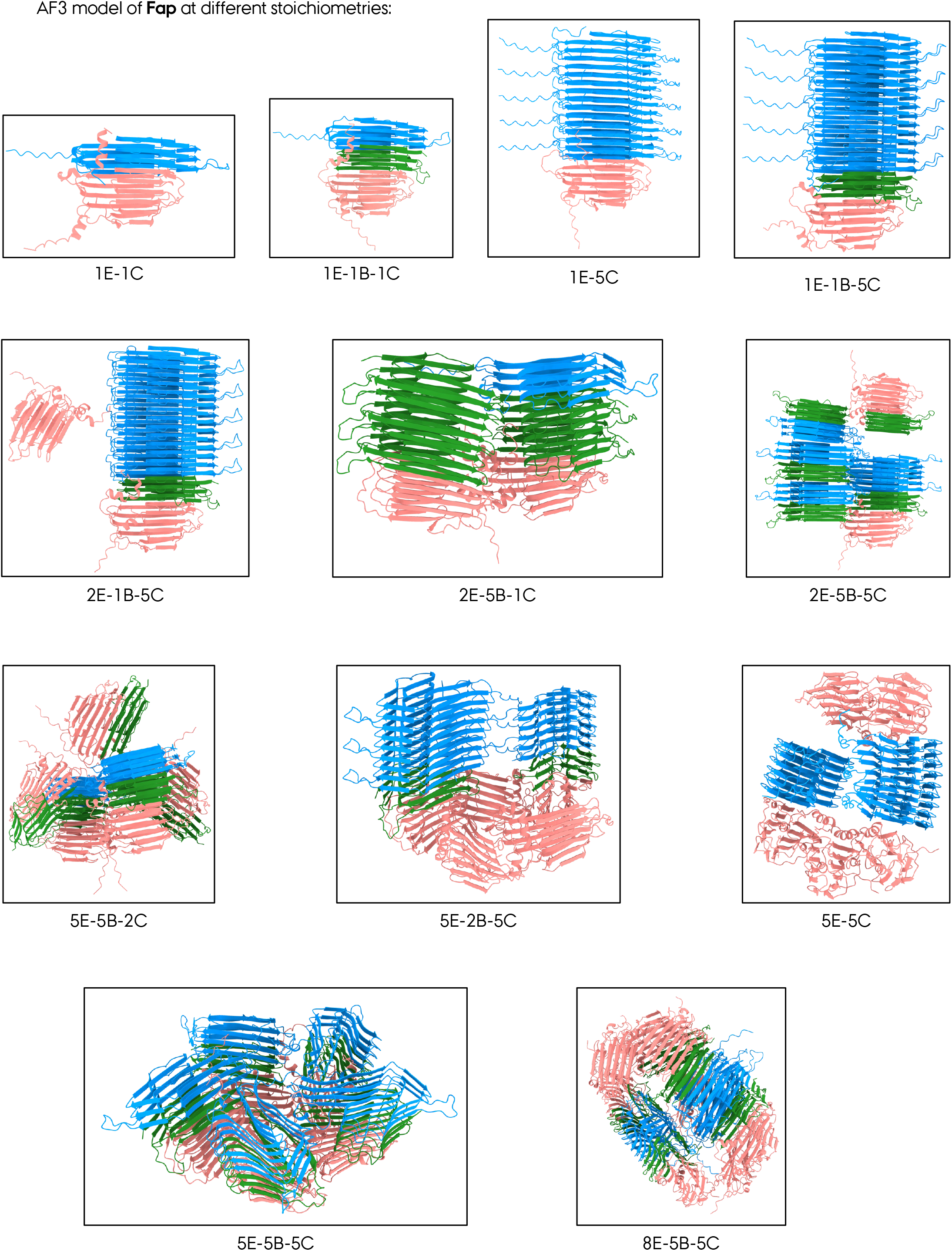
AF3 modeling of Fap proteins (FapB, FapC and FapE) at different biologically relevant conditions, stoichiometries and scenarios. FapE is pink, FapB is green and FapC is blue. The ratios of the three proteins are given below each box/condition. Protein sequences are used without signal sequences and modelled with random order of input at the AlphaFold Server (alphafoldserver.com).

**Fig. SI2:**
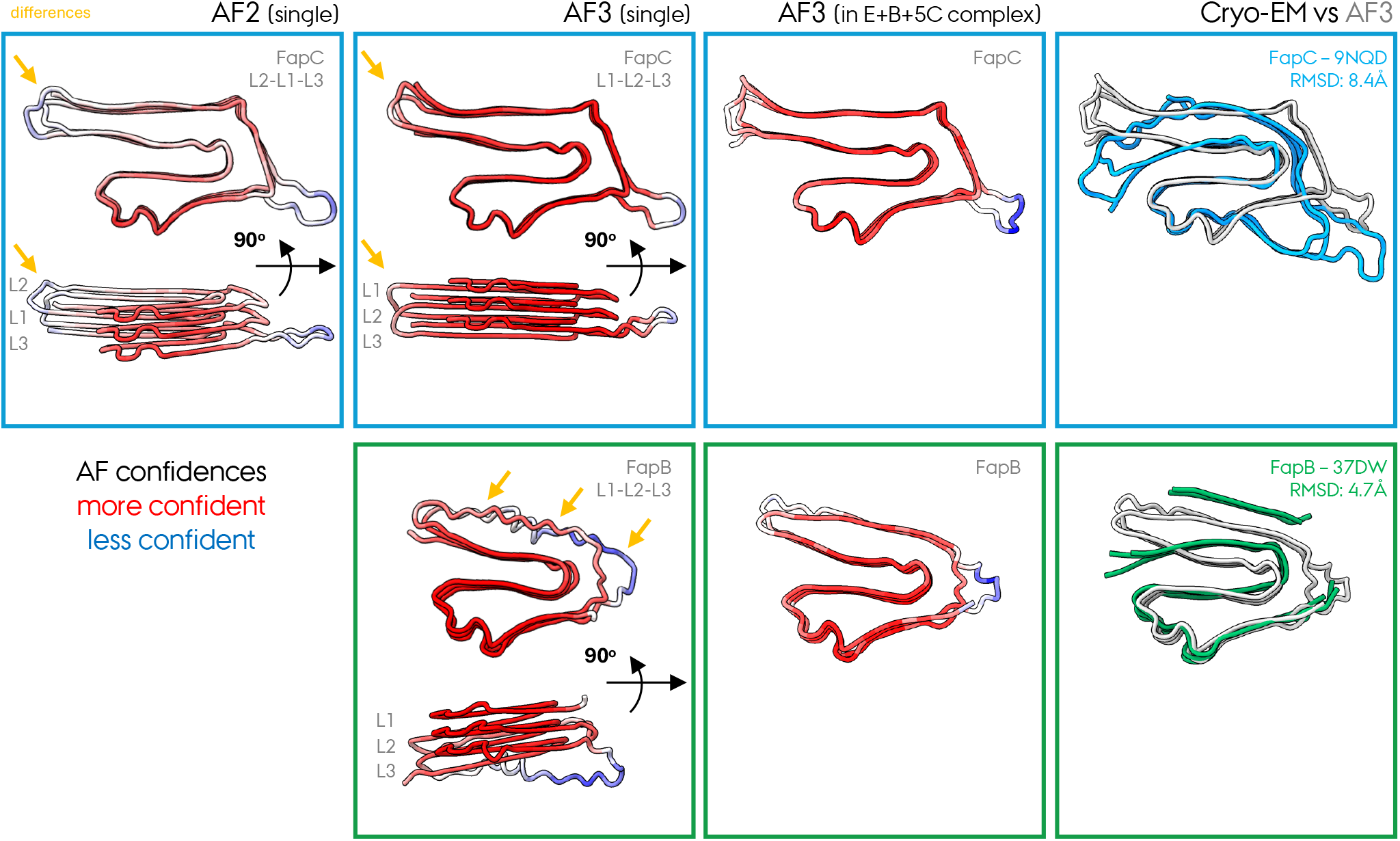
Comparison of FapB and FapC Cryo-EM structures with their AF2 and AF3 models. FapC is blue and FapB is green. The all-atom RMSD of the Cryo-EM and AF3 structures/models for FapB and FapC is given. For the AF2 and AF3 models of FapB and FapC, the confidence levels are visualized, where blue represents more confident areas and red represents less confident areas in the models. Different layer organization for FapC aF2 and AF3 models are visualized, where AF2 indicates an irregular/asymmetric L2-L1-L3 layer organization, whereas AF3 indicates a regular/symmetric L1-L2-L3 organization. Yellow arrows indicate differences in layer organization, confidence, and structural features, and are provided as a guide to the reader.

**Fig. SI3:**
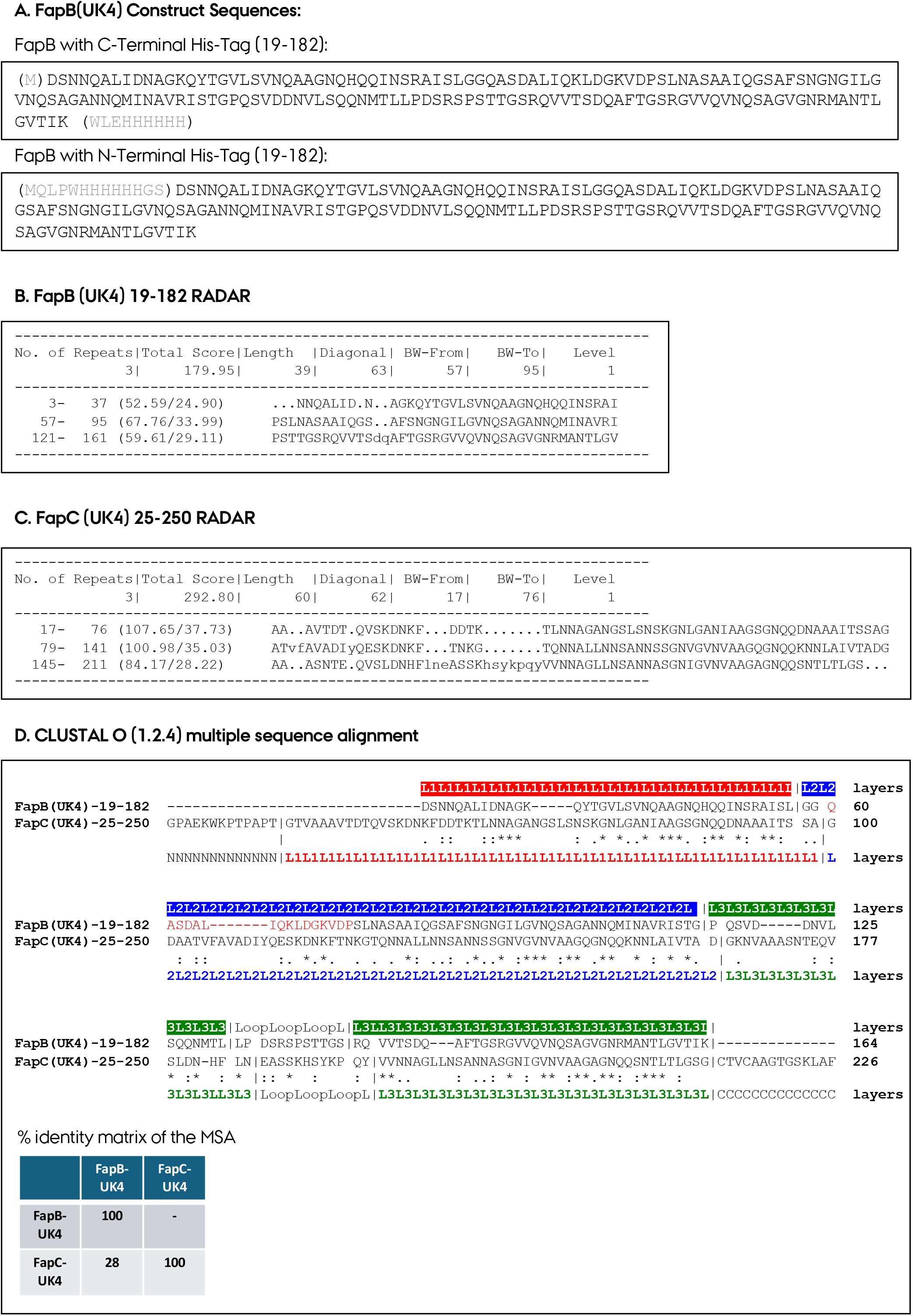
**A.** The sequences of the FapB and FapC protein constructs used throughout the study. The repeat prediction of **B.** UK4 FapB (19-182 residues) and **C.** UK4 FapC by RADAR. **D.** Multiple sequence alignment of FapB and FapC sequences without expression tags. The layer organization is given on top/bottom of the sequences for FapB and FapC, respectively. The % identity matrix is additionally shown resulting from the MSA.

**Fig. SI4:**
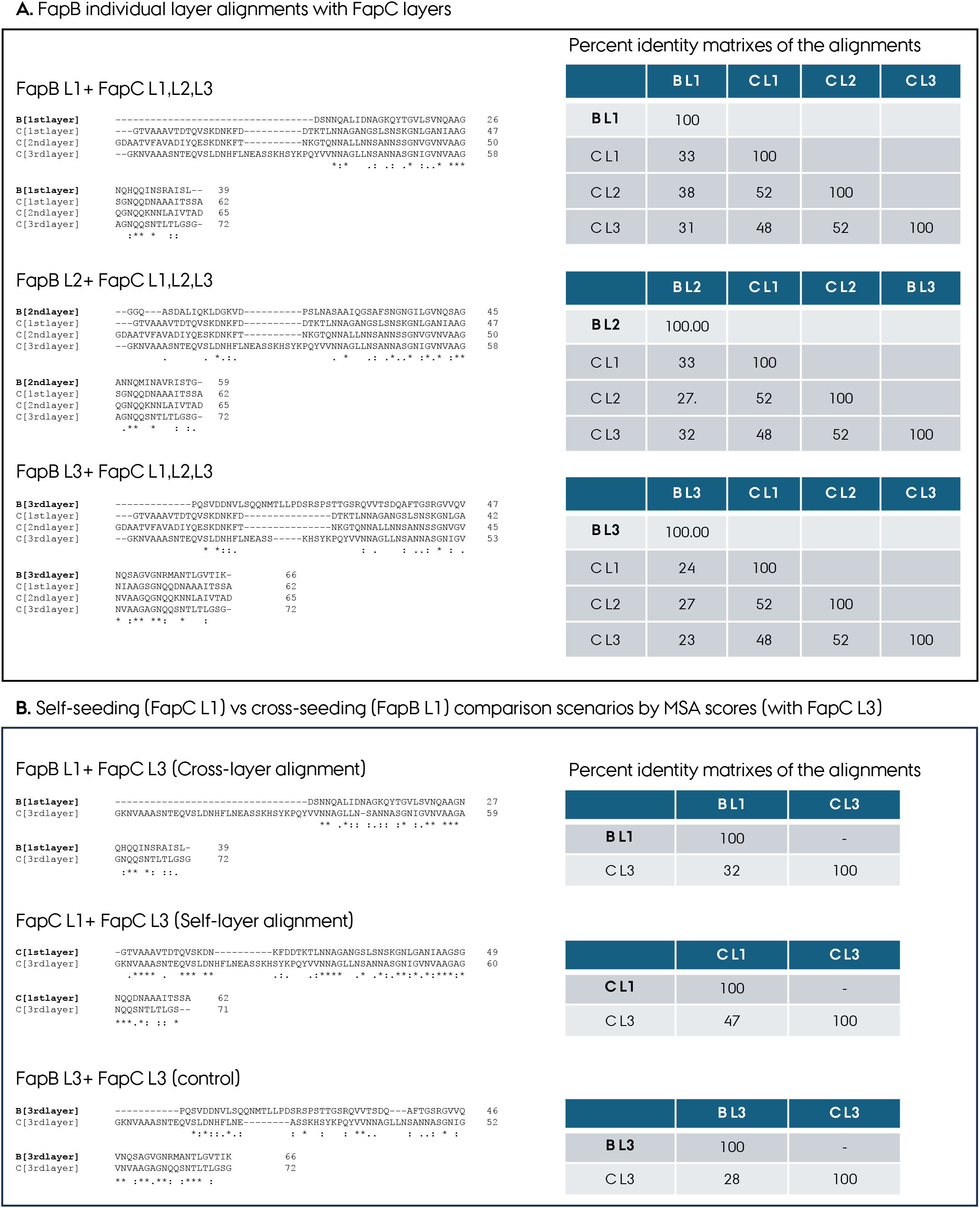
**A.** The multiple sequence alignment of FapB individual layers, L1-L3, with all three layers of FapC individually. The % identity matrix is additionally shown, resulting from the MSA. **B.** The self-seeding (FapC L1 with FapC L3) vs cross-seeding (FapB L1 with FapC L3) comparison scenarios by MSA scores and identity matrices. As a plausible control, the FapC L3 with FapC L3 scenario is also given.

**Fig. SI5:**
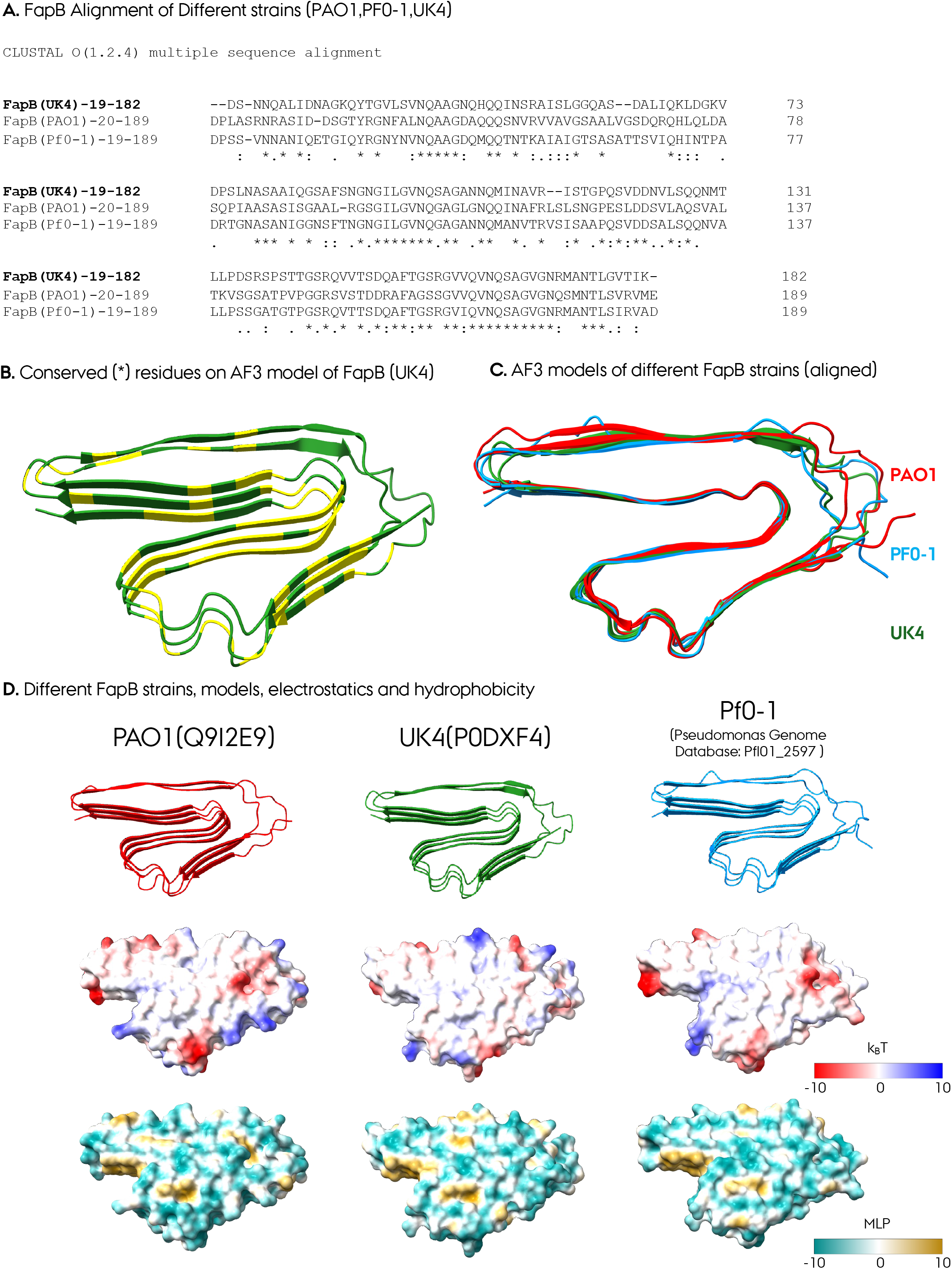
**A.** The multiple sequence alignment of FapB sequences from *Pseudomonas* UK4, PAO1, and Pf0-1 strains. **B.** The conserved residues (marked in yellow) originating from the MSA of these three strains shown in A are shown on the AF3 FapB UK4 model. **C.** AF3 models of these three strains represented as aligned to the UK4 model. **D.** The AF3 model of three *Pseudomonas* FapB strains, along with their electrostatic and hydrophobicity representations.

**Fig. SI6:**
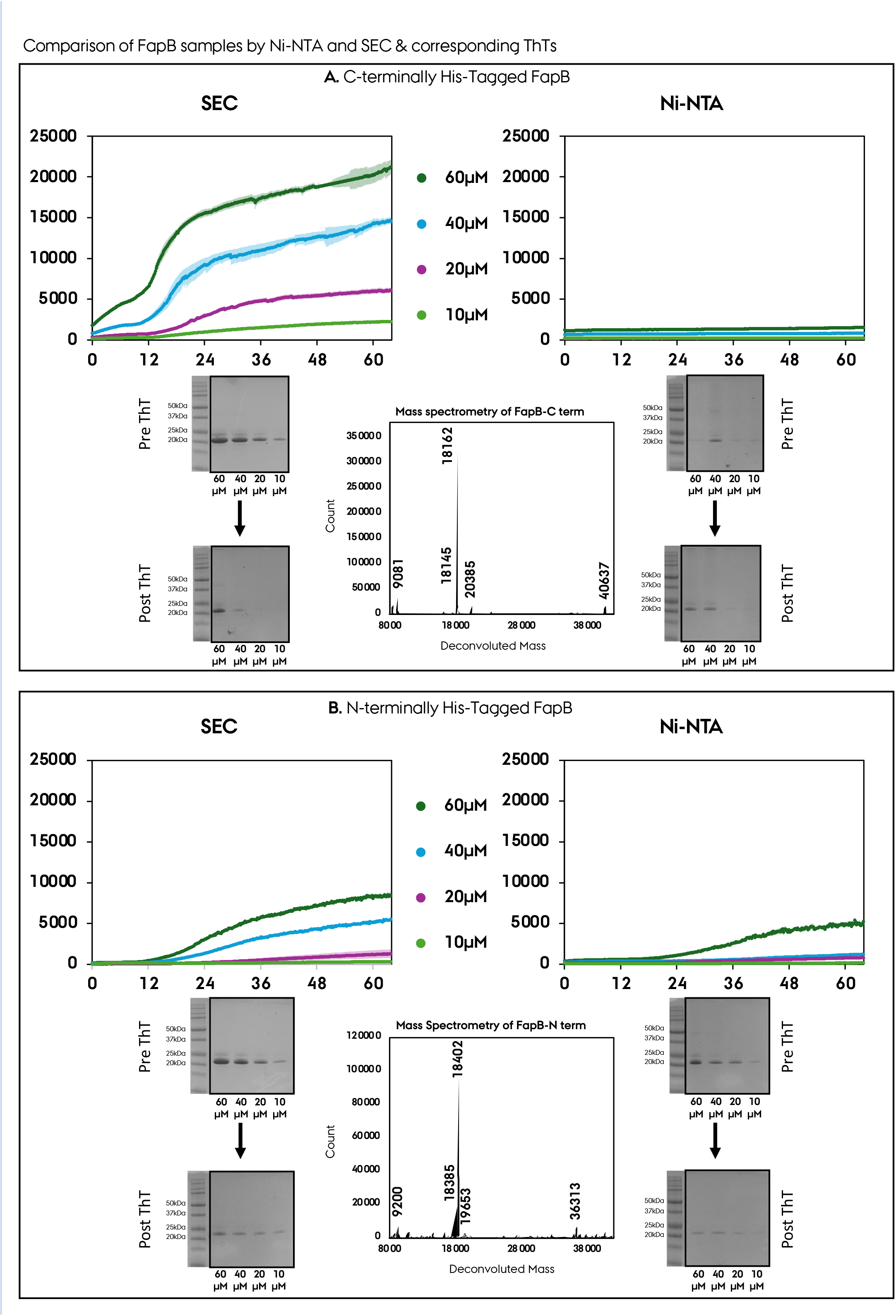
Comparison of **A.** FapB and **B.** FapC samples prepared and purified by Ni-NTA vs. SEC, along with their ThT curves, pre/post ThT SDS quantifications of monomer ratios, and the corresponding mass spectrometry (MS) results of the final purified sample.

**Fig. SI7:**
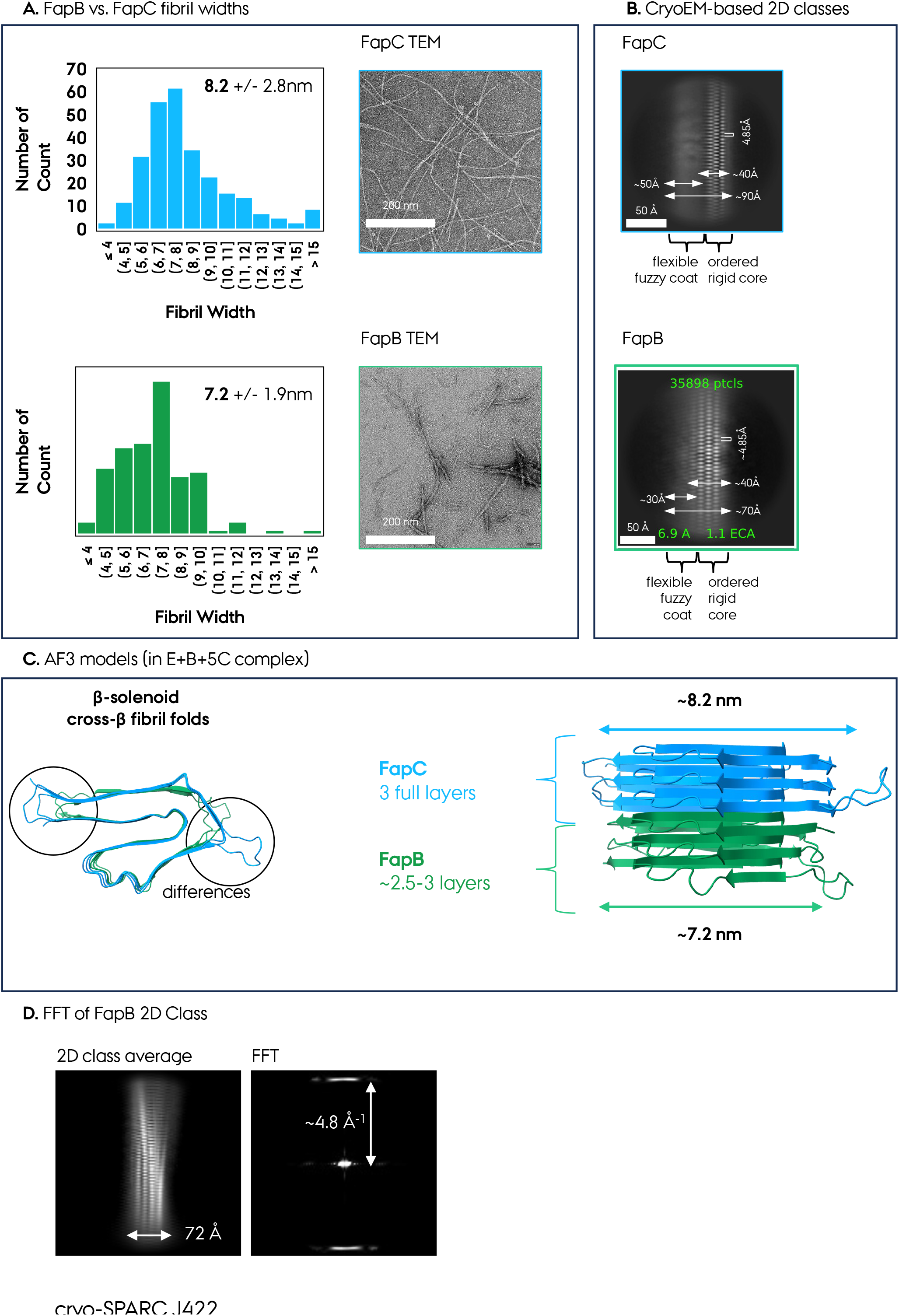
**A.** Representation of negative-staining transmission electron microscopy micrographs of FapB and FapC fibrils, along with their fibril width measurements and averages. **B.** The Cryo-EM 2D classes for FapB and FapC, along with the dimensions of the ordered fibril core and flexible fuzzy coat regions. **C.** Comparison of the AF3 models of FapB and FapC from top/side views, and the differences in their dimensions. **D.** The Fourier transform (FT) of the 2D class average for the FapB fibril with a dimension of ∼72 Å and the corresponding diffraction pattern marked at ∼4.8 Å.

**Fig. SI8:**
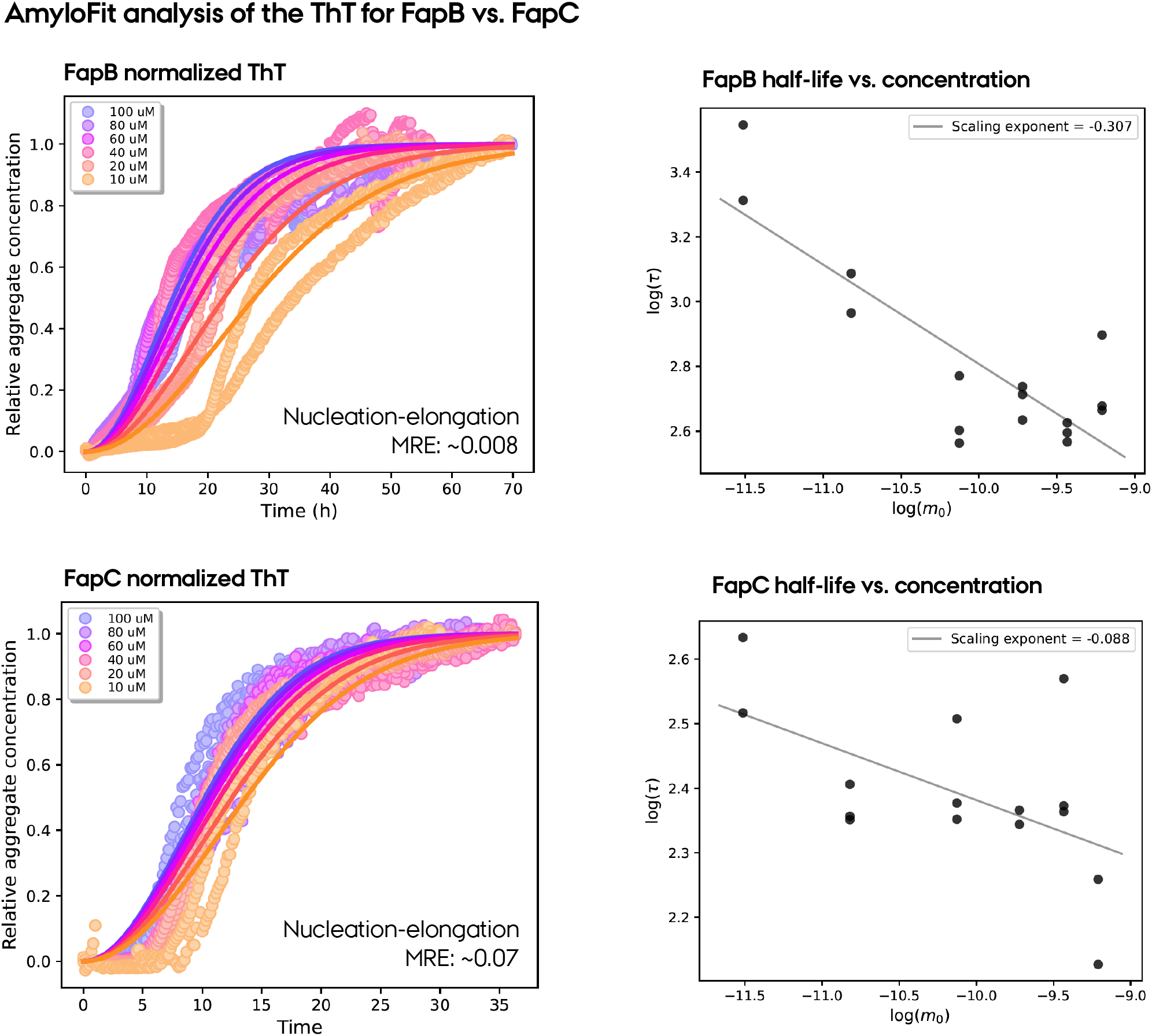
The fittings of the ThT curves of FapB and FapC performed at different concentrations between 10-100 μM. The mean residual error (MRE) values are given for each fit. Moreover, the corresponding half-life vs. log(concentration) plots are shown, which were utilized to determine the fibril formation mechanisms for FapB and FapC along with the substoichiometric self/cross-seeding assays.

**Fig. SI9:**
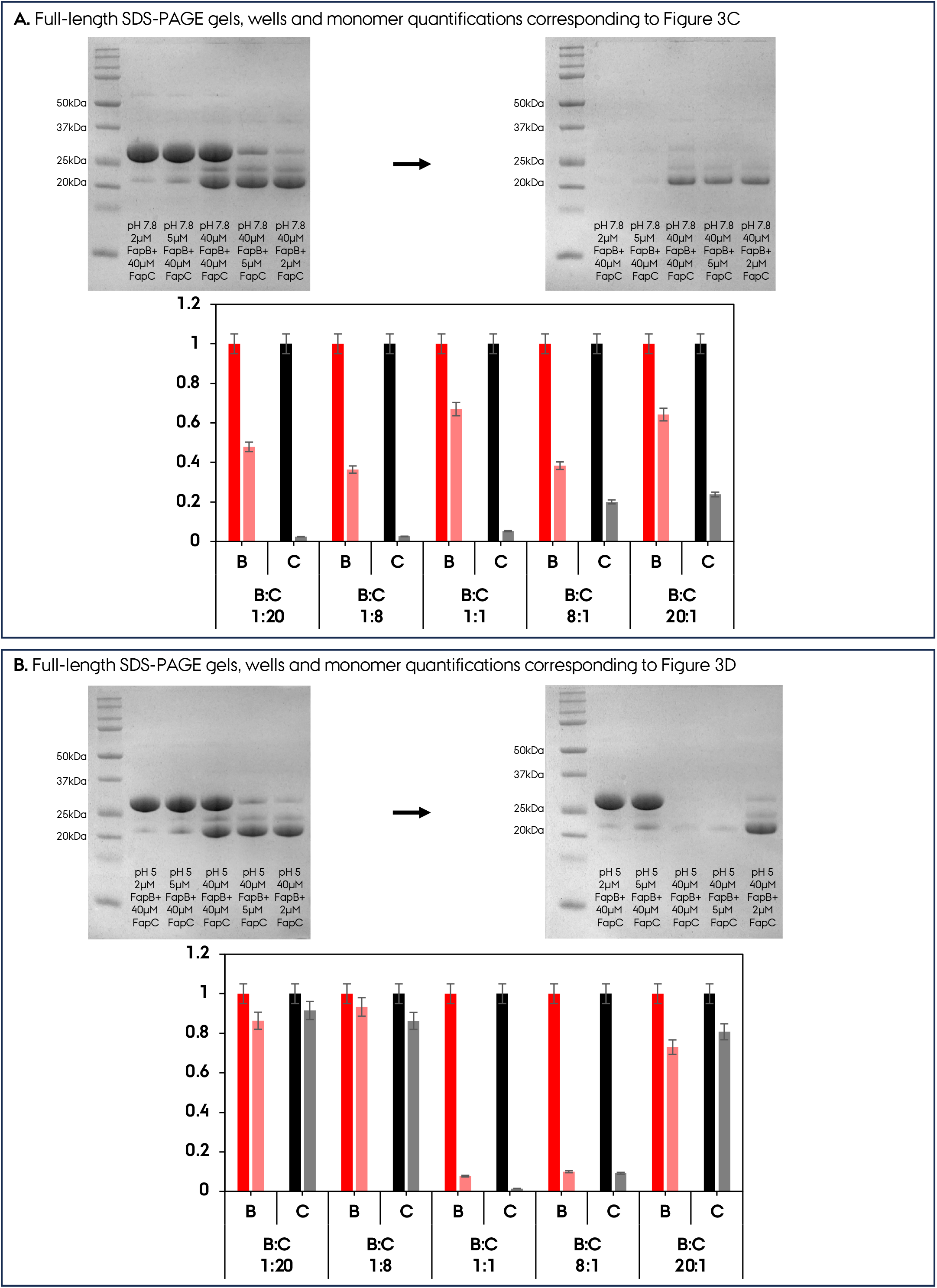
**A.** Full-length SDS-PAGE gels, wells and monomer quantifications corresponding to Figure 3C. **B.** Full-length SDS-PAGE gels, wells and monomer quantifications corresponding to Figure 3D.

**Fig. SI10:**
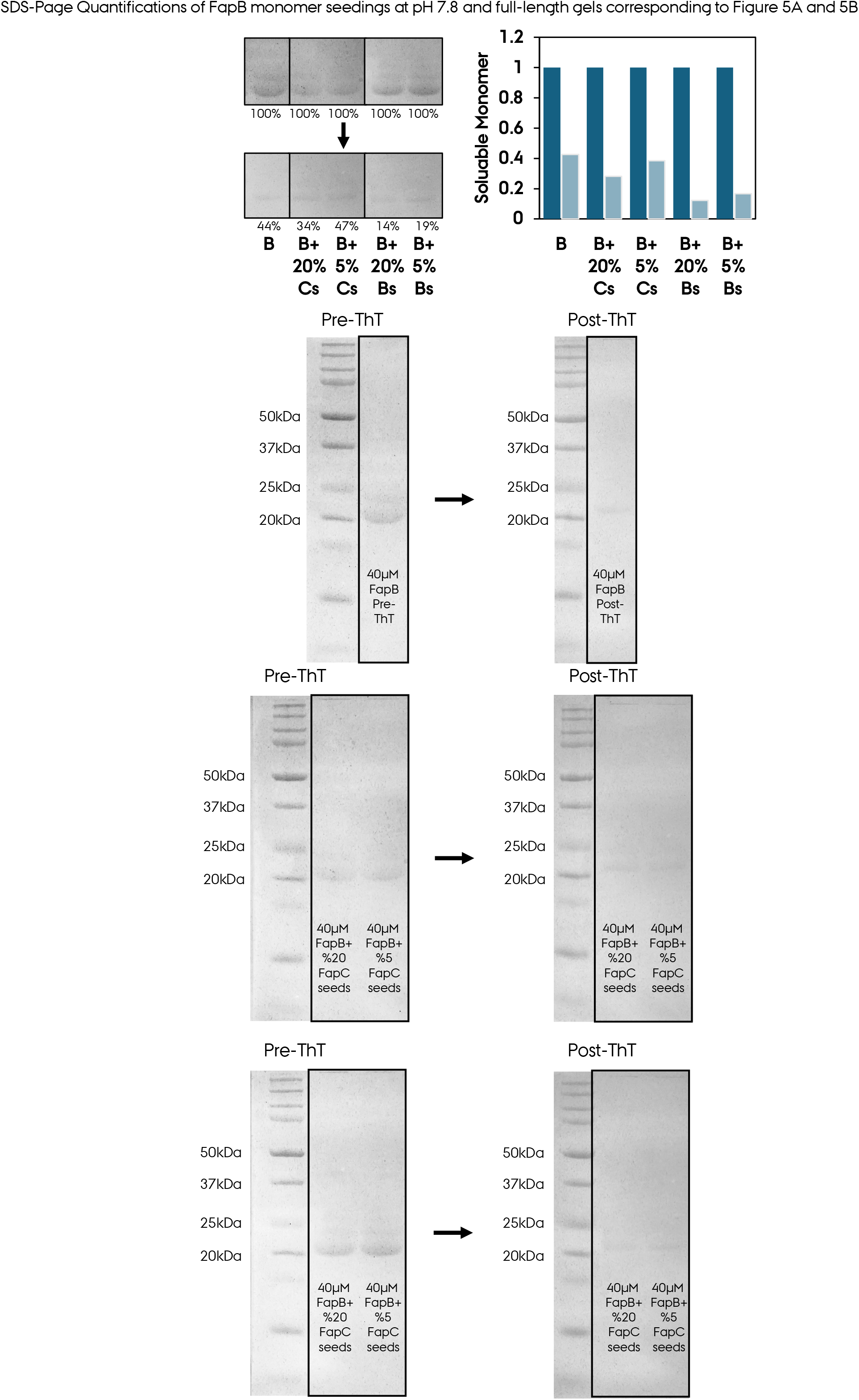
SDS-PAGE quantifications of FapB monomer seedings at pH 7.8 and full-length gels corresponding to Figure 5A and 5B, part 1.

**Fig. SI11:**
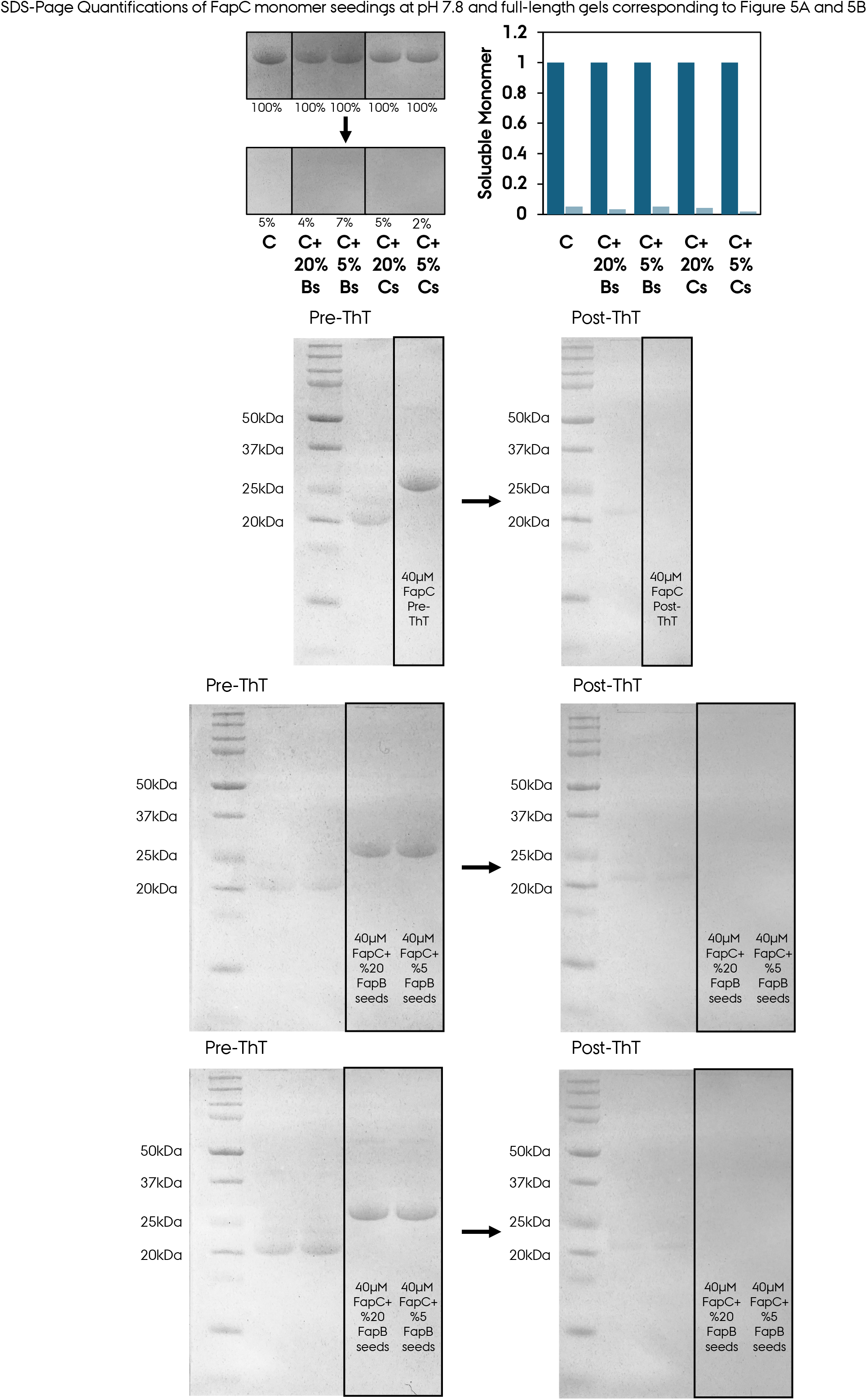
SDS-PAGE quantifications of FapC monomer seedings at pH 7.8 and full-length gels corresponding to Figure 5A and 5B, part 2.

**Fig. SI12:**
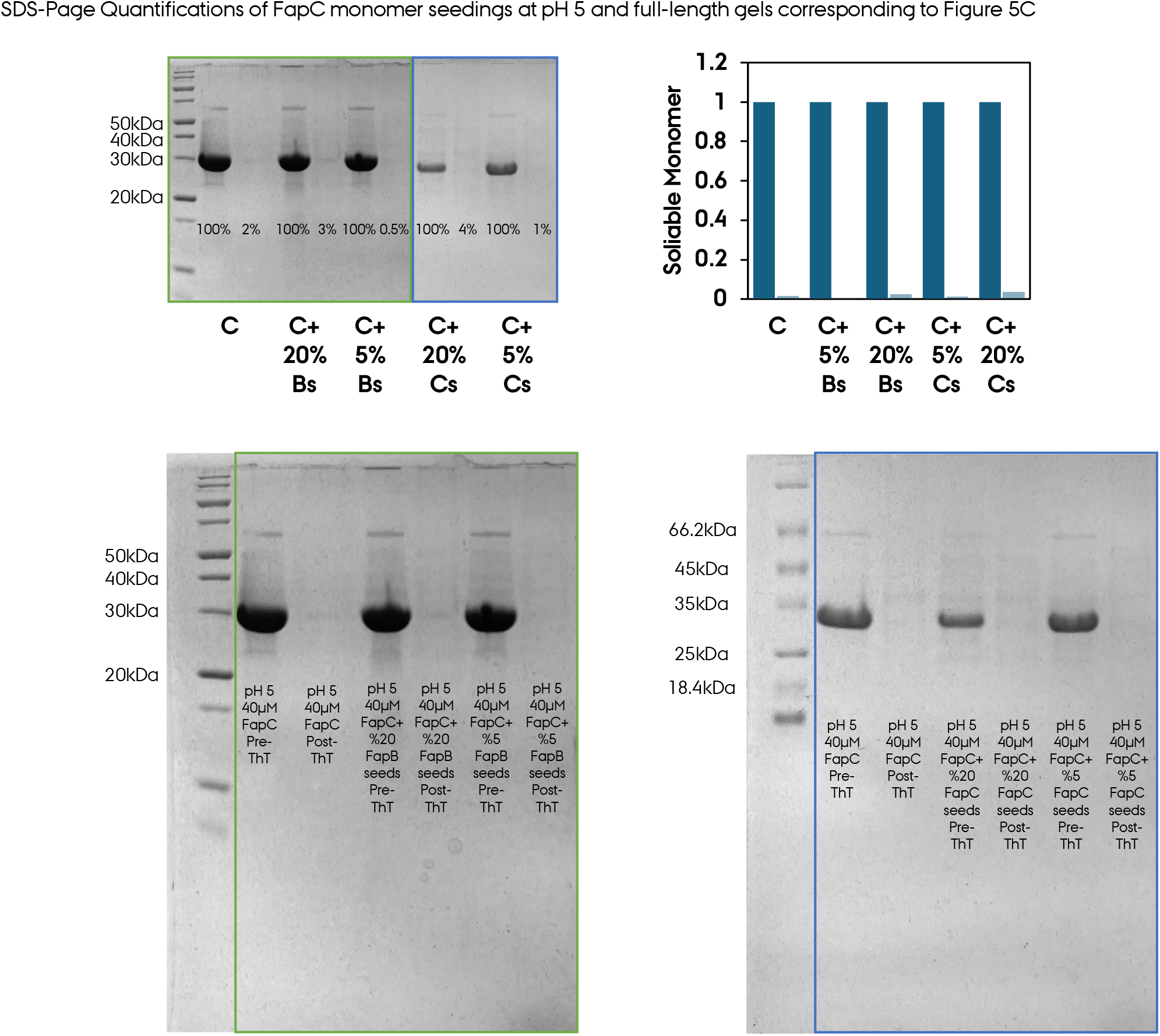
SDS-PAGE quantifications of FapC monomer seedings at pH 5 and full-length gels corresponding to Figure 5C.

**Fig. SI13:**
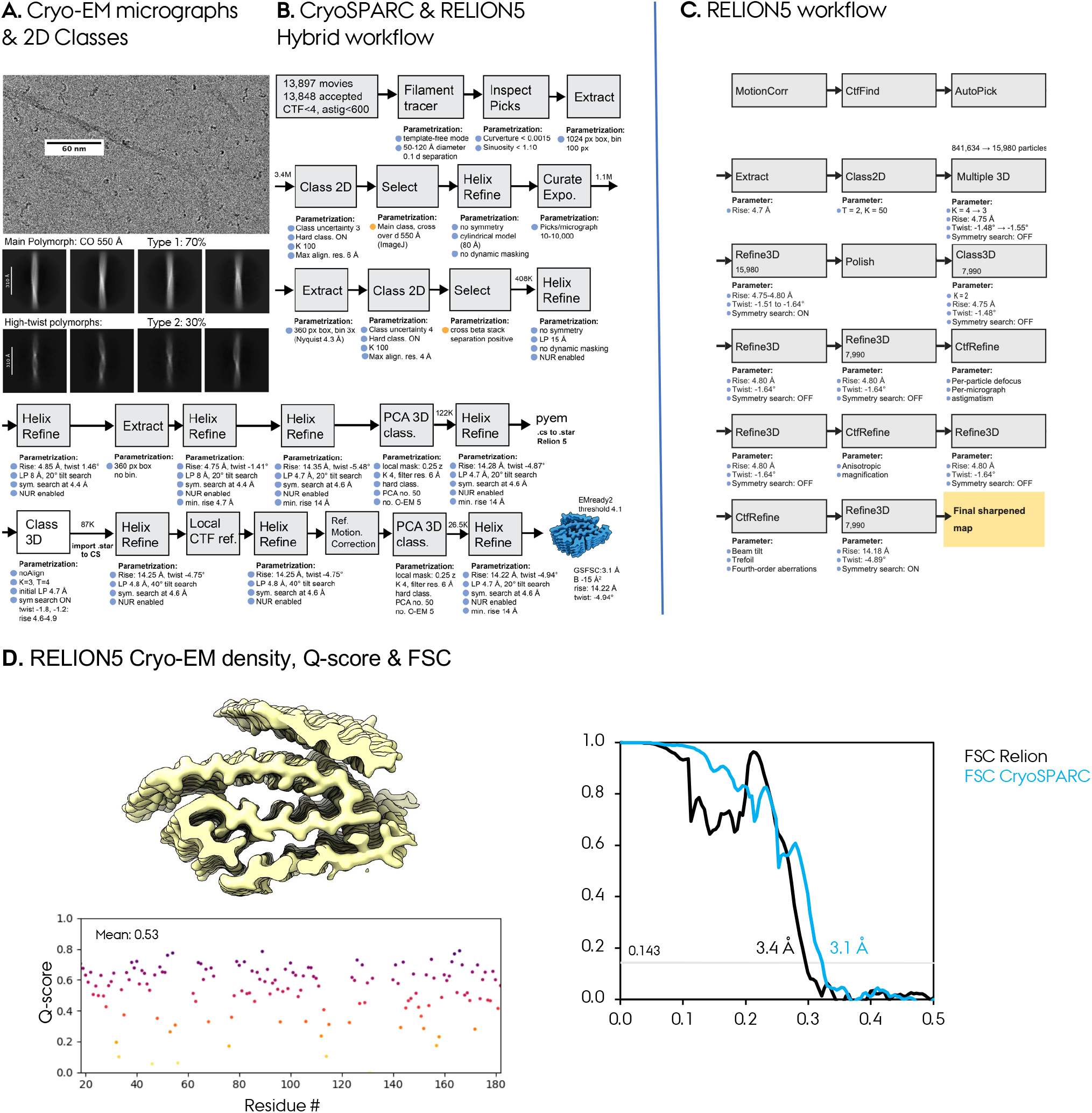
Helical Reconstruction of *Pseudomonas* FapB-UK4. **A.** Representative Cryo-EM micrographs for FapB fibrils (scale bar 60 nm), along with the resulting 2D Class averages corresponding to two different types of FapB fibrils/polymorphs. 2D class averages of the main polymorph with a cross-over (CO) distance of 550 Å and other non-resolved high-twist polymorphs; scale bars 310 Å. Flow chart representing the protocol utilized to determine FapB Cryo-EM structures by **B.** CryoSPARC alone and also the CryoSPARC + RELION5 hybrid approach (blue parameter points are automatable; orange points require manual intervention); **C.** RELION5 approach. **D.** The final RELION -based Cryo-EM density of FapB, from the major polymorph Type-1 with 550 Å CO distance. The Q-scores for the map-model fitting are given per residue. For the validation of FapB Cryo-EM density from processing in cryoSPARC, a Gold-Standard Fourier-Shell Correlation (GS-FSC) reporting the overall resolution at FSC cutoff 0.143 is given (3.1 Å for the CryoSPARC hybrid workflow and 3.4 Å for the RELION5 workflow).

**Fig. SI14:**
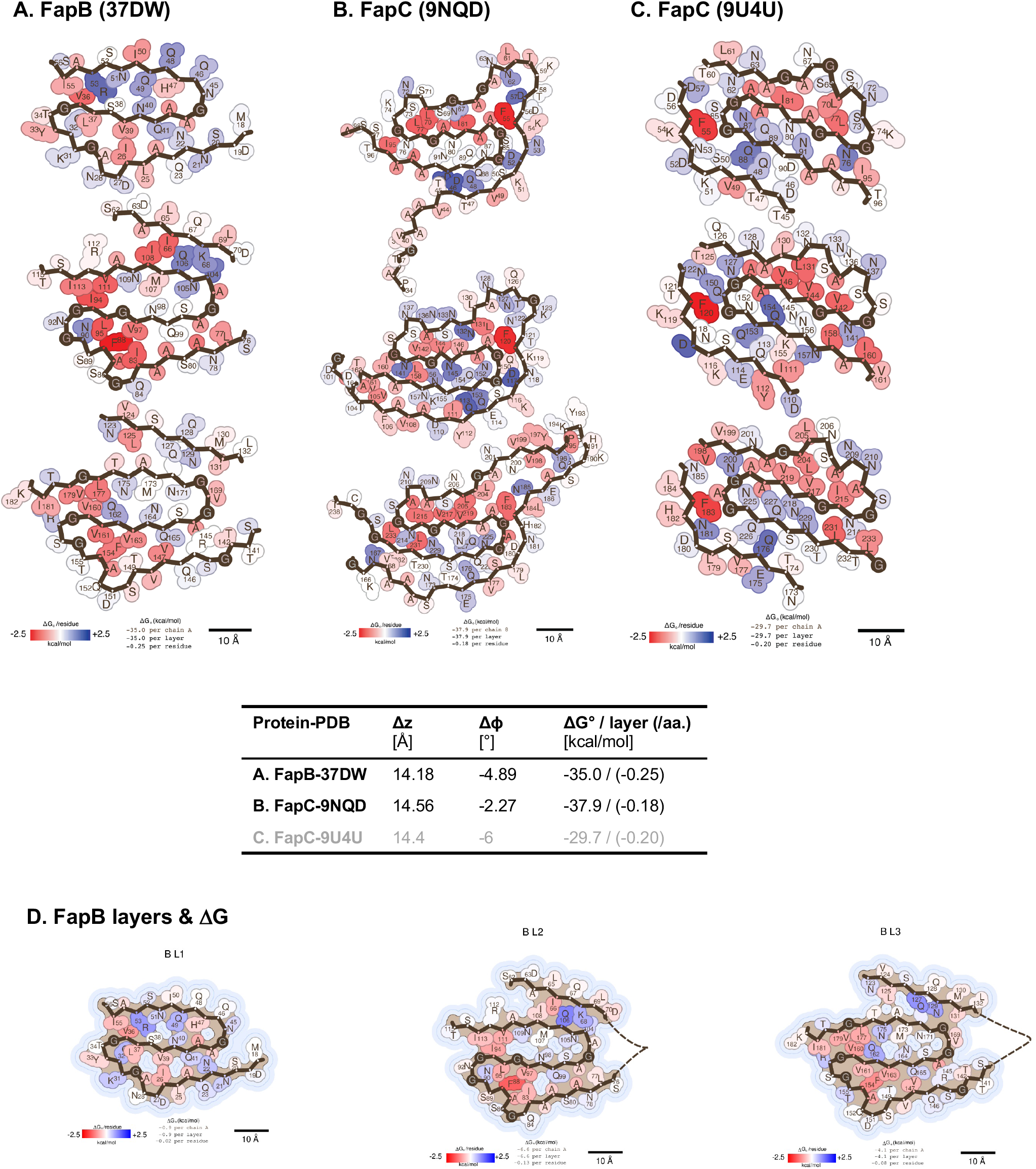
Per-layer and per-residue amyloid solvation energies (**Δ**G°) from Amyloid Illustrator for **A.** FapB Cryo-EM structure presented here (37DW), **B.** FapC (9NQD and **C.** FapC (9U4U). Red: stabilizing; blue: destabilizing residue. The combined helical parameters and solvation-energy values per layer and per residue for the current FapB and two FapC structures are also given in a table. **D.** The amyloid solvation energies (**Δ**G°) from Amyloid Illustrator for three FapB layers L1-L3.

**Fig. SI15:**
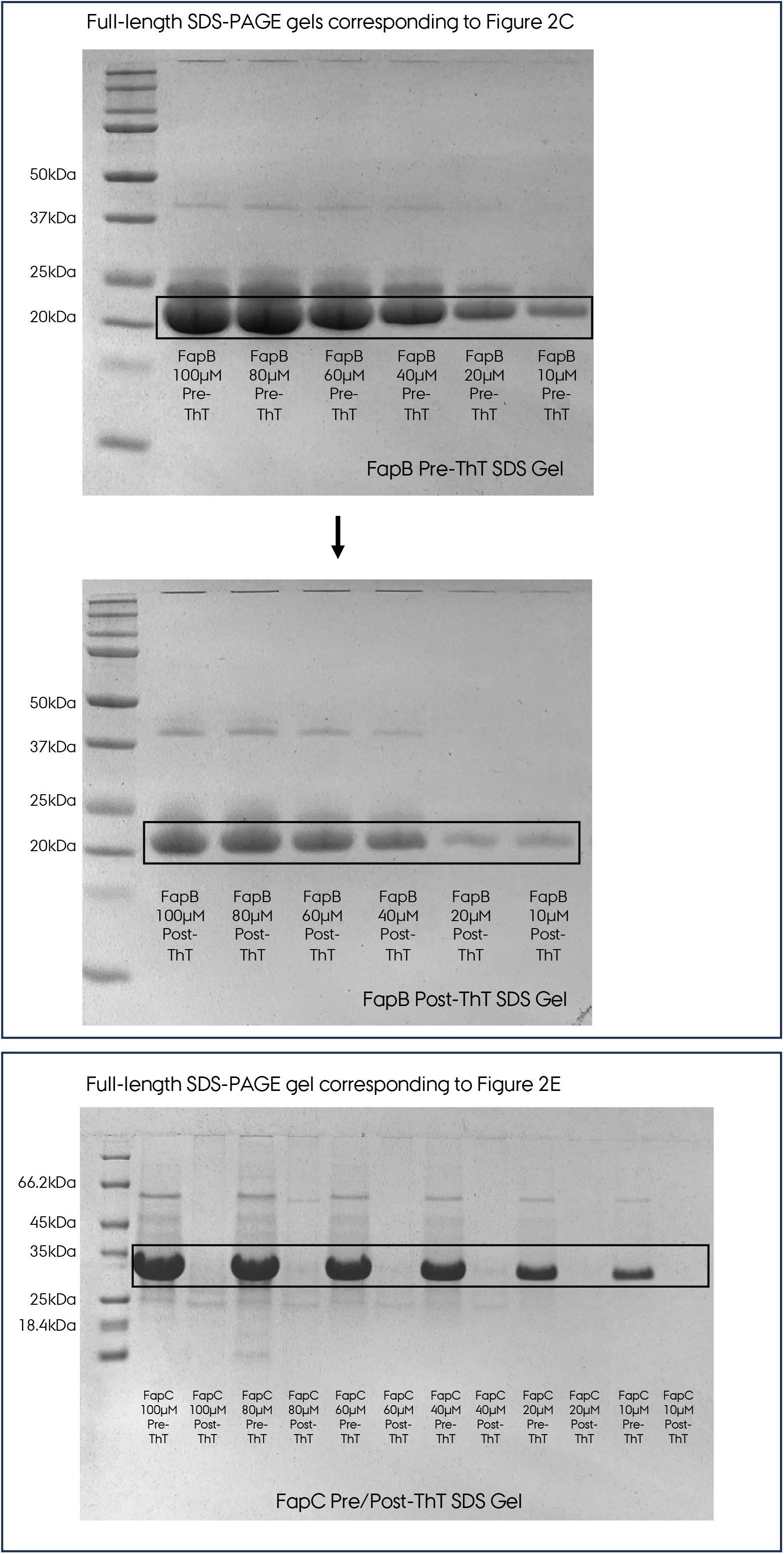
Full-length SDS-PAGE gels corresponding to Figure 2C and Figure 2E.

**Fig. SI16:**
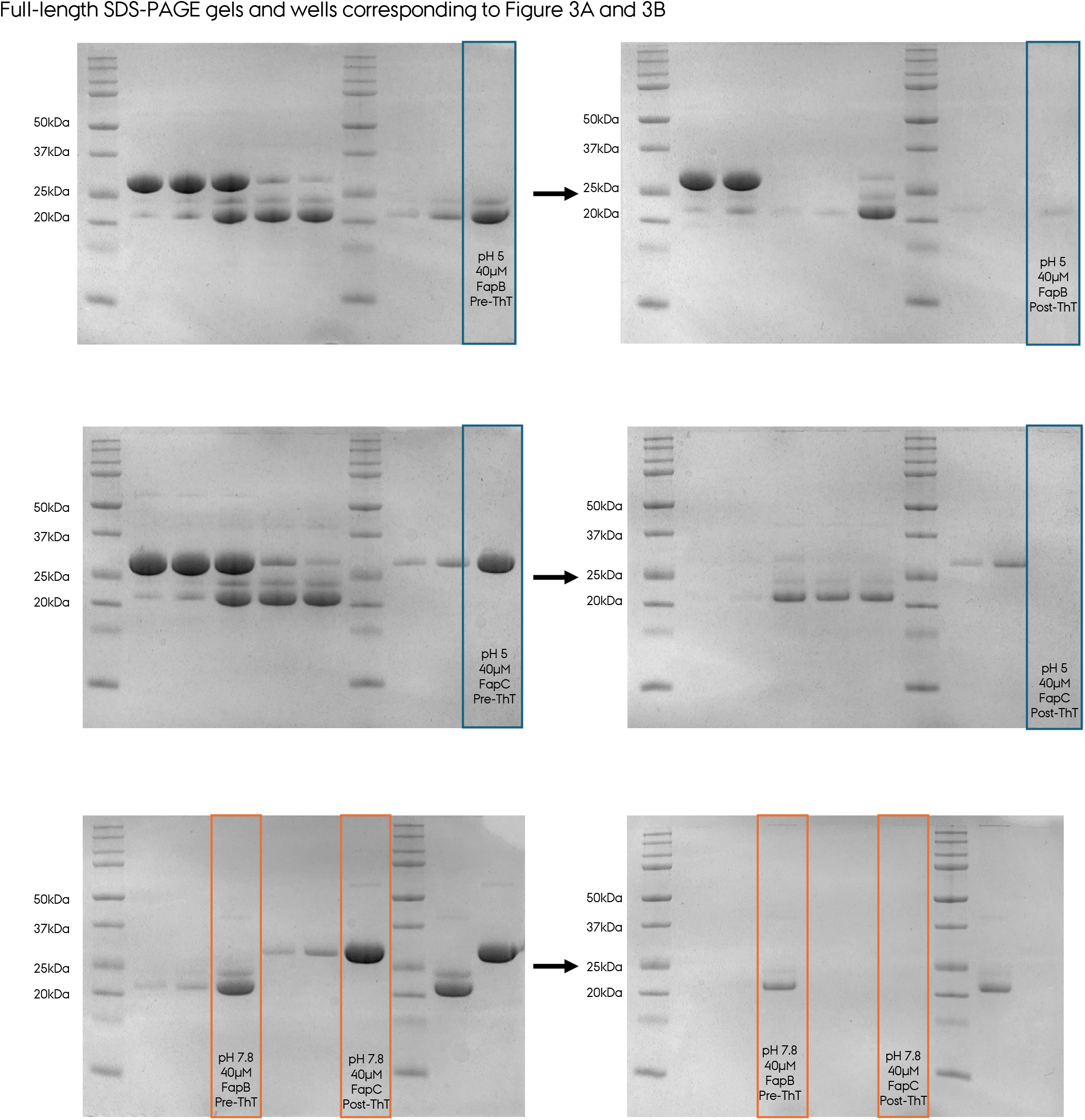
Full-length SDS-PAGE gels and wells corresponding to Figure 3A and 3B.

**Table S1.** Cryo-EM data collection and image processing of amyloids. (NA, not applicable. -, data not available).

|  | **FapB-UK4 (Uniprot: P0DXF4)** |
| --- | --- |
| EMDB ID | EMD-78102 |
| PDB ID | 37DW |
| **Data Collection** |  |
| Microscope | TFS Krios |
| Camera | Falcon 4i |
| Magnification (nominal) | 165,000 |
| Voltage (keV) | 300 |
| Total dose (e<sup>-</sup>/Å<sup>2</sup>) | 50 |
| EER Fractions | 18 |
| Pixel size (Å/px) | 0.72 |
| Defocus range (μm) | -0.5 to -2.5 |
| Recorded movies | 13,897 |
| **Image Processing** |  |
| Processing software | RELION5 |
| Final particle images | ~8,000 |
| Symmetry imposed | C1 |
| Rise (Å) | 14.18 |
| Twist (°) | -4.89 |
| Res. (FSC 0.143) (Å) | 3.4 |
| Local res. range (Å) | 5.07 to 3.02 |
| **Model Refinement** |  |
| Initial model | AlphaFold3 |
| MolProbity score | 1.26 |
| Clash Score | 4.92 |
| **Composition** |  |
| Chains | 3 |
| Atoms | 7,518 |
| Protein residues | 522 |
| **Bonds (R.M.S.D.)** |  |
| Length (Å) | 0.013 |
| Angles (°) | 2.313 |
| **B-factors (min/max/mean)** |  |
| Protein residues | 81.48/196.85/118.14 |
| **Ramachandran plot (%)** |  |
| Favored | 98.53 |
| Allowed | 1.47 |
| Outliers | 0.00 |
| Rotamer outliers (%) | 0.00 |
| Q-score | 0.52 |

